# Dichotomy between extracellular signatures of active dendritic chemical synapses and gap junctions

**DOI:** 10.1101/2024.07.04.602060

**Authors:** Richa Sirmaur, Rishikesh Narayanan

## Abstract

Local field potentials (LFPs) are compound signals that represent the dynamic flow of information across the brain, which have been historically associated with chemical synaptic inputs. How do gap junctional inputs onto active compartments shape LFPs? We developed methodology to record extracellular potentials associated with different patterns of gap junctional inputs onto conductance-based models. We found that synchronous inputs through chemical synapses yielded a negative deflection in proximal extracellular electrodes, whereas those onto gap junctions manifested a positive deflection. Importantly, we observed extracellular dipoles only when inputs arrived through chemical synapses, but not with gap junctions. Remarkably, hyperpolarization-activation cyclic nucleotide-gated channels, which typically conduct inward currents, mediated outward currents triggered by the fast voltage transition caused by synchronous inputs. With rhythmic inputs at different frequencies arriving through gap junctions, we found strong suppression of LFP power at higher frequencies as well as frequency-dependent differences in the spike phase associated with the LFP, when compared to respective chemical synaptic counterparts. All observed differences in LFP were mediated by the relative dominance of synaptic currents *vs*. voltage-driven transmembrane currents with chemical synapses *vs*. gap junctions, respectively. Our analyses unveil a hitherto unknown role for active dendritic gap junctions in shaping extracellular potentials.

## INTRODUCTION

Perpetual ionic movement across membranes and within cells play core physiological roles in the brain. Such ionic movements result in electrical potentials that can be recorded from the extracellular milieu, and are referred to as extracellular field potentials (EFPs). Different types of extracellular potentials, including electroencephalogram, electrocorticogram, and potentials from cell-proximal electrodes, have offered critical insights about brain physiology and have also proved useful in devising brain-computer interfaces (Buzsaki, 2004, 2006; Buzsaki et al., 2012; Einevoll et al., 2013; Obien et al., 2014; Lebedev and Nicolelis, 2017; Pesaran et al., 2018; Pandarinath and Bensmaia, 2022; Halnes et al., 2024). The high-frequency components of cell-proximal extracellular recordings provide spiking information about adjacent neurons. The low-frequency components of these extracellular potentials, referred to as local field potentials (LFP), were traditionally thought to be reflective of currents through synaptic receptors located on different postsynaptic compartments. This synapse-centric view of LFPs was considerably revised after the discovery of active dendrites, to include currents through different ion channels into LFP analyses (Buzsaki et al., 2012; Schomburg et al., 2012; Reimann et al., 2013; Anastassiou et al., 2015; Sinha and Narayanan, 2015; Ness et al., 2016, 2018; Sinha and Narayanan, 2022). However, an important caveat in most LFP analyses is that they are limited to scenarios where synaptic inputs are assumed to be exclusively arriving through chemical synapses that regulate extracellular potentials through transmembrane receptor currents.

Cells in the brain communicate with each other through not just chemical synapses (Dale, 1934; Loewi, 1945; Bullock, 1951), but also through gap junctions (Curtis and Travis, 1951; Furshpan and Potter, 1957; Watanabe, 1958). Connectivity through gap junctions manifests as protoplasmic extensions through specialized molecules implementing a physical continuity between two cells that they connect. Chemical synapses and gap junctions express across different compartments between various cell types across the brain, with mixed connectivity known to manifest between certain cell types (Connors and Long, 2004; Andersen et al., 2006; Pereda, 2014; Connors, 2017; Nagy et al., 2018; Verkhratsky and Nedergaard, 2018).

A fundamental caveat in LFP analyses, therefore, is the omission of inputs arriving through gap junctions, especially onto structures that express an abundance of active mechanisms that mediate transmembrane currents. Incorporation of gap junctional inputs into LFP analyses becomes essential especially because of the several physiological roles played by gap junctions, including in bolstering burst firing and synchronization in different oscillatory bands (Christie et al., 1989; Kepler et al., 1990; Sherman and Rinzel, 1992; Draguhn et al., 1998; Skinner et al., 1999; Alvarez et al., 2002; Traub et al., 2003b; Bennett and Zukin, 2004; Kopell and Ermentrout, 2004; Traub et al., 2011; Connors, 2017). The extracellular potential associated with chemical synapses on a passive membrane would reflect the transmembrane synaptic current as well as the capacitive and leak currents driven by the voltage response to the synaptic current. However, for chemical synapses on an active structure, currents driven by the voltage response would also include all the voltage– and calcium-gated ion channels on the membrane. Unlike chemical synapses, gap junctions do not manifest a transmembrane synaptic current. When postsynaptic structures were considered to be passive, gap junctional impact on field potentials was considered to be indirect through modulation of neural excitability (Halnes et al., 2024). This view, however, needs marked revision to account for the active nature of dendrites (Johnston and Narayanan, 2008; Major et al., 2013; Stuart and Spruston, 2015) and other compartments (Traub and Bibbig, 2000; Traub et al., 2012; Verkhratsky and Nedergaard, 2018) that receive gap junctional inputs.

Analyses of extracellular potentials with gap junctions on active structures must explicitly account for the several transmembrane currents, including the capacitive, leak, and all active currents driven by voltage response to the gap junctional current, towards addressing several important questions. How do similar kinds of inputs arriving through chemical synapses *vs*. gap junctions impact extracellular potentials? What contributions do active structures receiving chemical synapses *vs*. gap junctions make to extracellular potentials? How does the absence of receptor transmembrane current with gap junctional inputs alter potentials across different cell-proximal extracellular electrode locations? Are there differences between the spectral power of extracellular potentials associated with oscillatory inputs arriving onto active or passive structures through chemical synapses *vs*. gap junctions? How does the chemical *vs*. electrical nature of synaptic input alter the relationship between extracellular voltages and intracellular voltages/spikes?

In this study, we employed morphologically realistic neuronal models in different configurations to address these questions about extracellular signatures of chemical synaptic *vs*. gap junctional inputs impinging on active or passive structures. Given the complexity arising from the concurrent activity of chemical synapses, gap junctions, and active dendritic conductances across multiple neuronal populations, experimentally isolating the contributions of individual components to extracellular potentials remains highly challenging. To address this limitation, we employed a computational modeling approach, which provides a quantitative framework for systematically dissecting the distinct roles of specific cellular and synaptic elements. This strategy is consistent with previous studies that have successfully used computational methods to elucidate the contributions of active dendritic mechanisms to LFPs (Reimann et al., 2013; Sinha and Narayanan, 2015; Ness et al., 2018; Sinha and Narayanan, 2022; Halnes et al., 2024) or spiking contributions to LFPs (Gold et al., 2006; Buzsaki et al., 2012; Schomburg et al., 2012). In addition, experimentally isolating the contribution of gap junctions is complicated by non-specific effects of available pharmacological modulators targeting these connections (Rouach et al., 2003; Behrens et al., 2011). Most genetic knockouts of gap junctional proteins are either lethal or trigger functional compensatory mechanisms (Lo, 1999; Bedner et al., 2012), thereby rendering causal attribution of specific gap junctional contributions infeasible with currently available experimental approaches. Consequently, biophysically and morphologically detailed computational modeling provides a crucial means to evaluate the impact of individual neuronal components on extracellular field potentials (Einevoll et al., 2019; Halnes et al., 2024).

Our analyses unveiled striking differences in polarities, spatiotemporal patterns, spectral signatures, and intracellular voltage relationships of extracellular potentials associated with inputs impinging on active dendrites through chemical synapses *vs*. gap junctions. We found these differences to be mediated by the differential dominance of receptor *vs*. active transmembrane currents associated with chemical synaptic *vs*. gap junctional inputs, respectively. The key distinctions in extracellular signatures of gap junctional inputs onto active structures emphasize the inadequacies associated with analyses that account solely for chemical synapses, strongly warranting the need to account for gap junctions in the analyses of extracellular potentials.

## RESULTS

### Synchronous inputs: Contrasting patterns of extracellular signatures associated with active dendritic chemical synapses *vs*. gap junctions

First, we assessed the impact of synchronous excitatory inputs impinging on active or passive basal dendrites, either through chemical synapses or gap junctions, on extracellular potentials. To do this, several synapses or junctions that were randomly placed at different locations along the dendritic tree were synchronously activated (Supplementary Fig. S1). Intracellularly, these synchronous inputs yielded a somatic action potential irrespective of whether they arrived through chemical synapses or gap junctions. In striking contrast, extracellular signatures manifested a strong dependence on whether these synchronous inputs arrived through chemical synapses (Fig. 1*A*) or gap junctions (Fig. 1*B*). Specifically, the extracellular potential manifested as a sink (a negative deflection) at proximal and distal basal locations with synchronous excitatory chemical synapses (Fig. 1*A*). However, when synchronous gap junctional connectivity provided the excitatory inputs, extracellular potentials manifested a source (a positive deflection) at proximal and distal basal locations (Fig. 1*B*). The amplitudes of extracellular potentials, across all electrode locations spanning the basal tree, were comparable for analyses with active or passive dendrites (with the soma remaining active for either scenario) when synchronous stimulation arrived through chemical synapses (Fig. 1*C*). Remarkably however, the amplitudes of extracellular potentials showed significant reductions between active and passive cases, across all electrode locations spanning the basal tree, when synchronous inputs arrived through gap junctions (Fig. 1*D*). These observations held even when the number of synaptic or junctional contacts were reduced (Supplementary Fig. S2), and together point to dominant contributions of transmembrane currents from active dendrites to extracellular field potentials associated with synchronous gap junctional inputs (Fig. 1*D*). The striking reduction in the amplitudes, spanning orders of magnitude, of extracellular potentials with passive dendrites (Fig. 1*D*) explains why gap junctional contributions to LFPs have been historically neglected.

**Figure 1.**
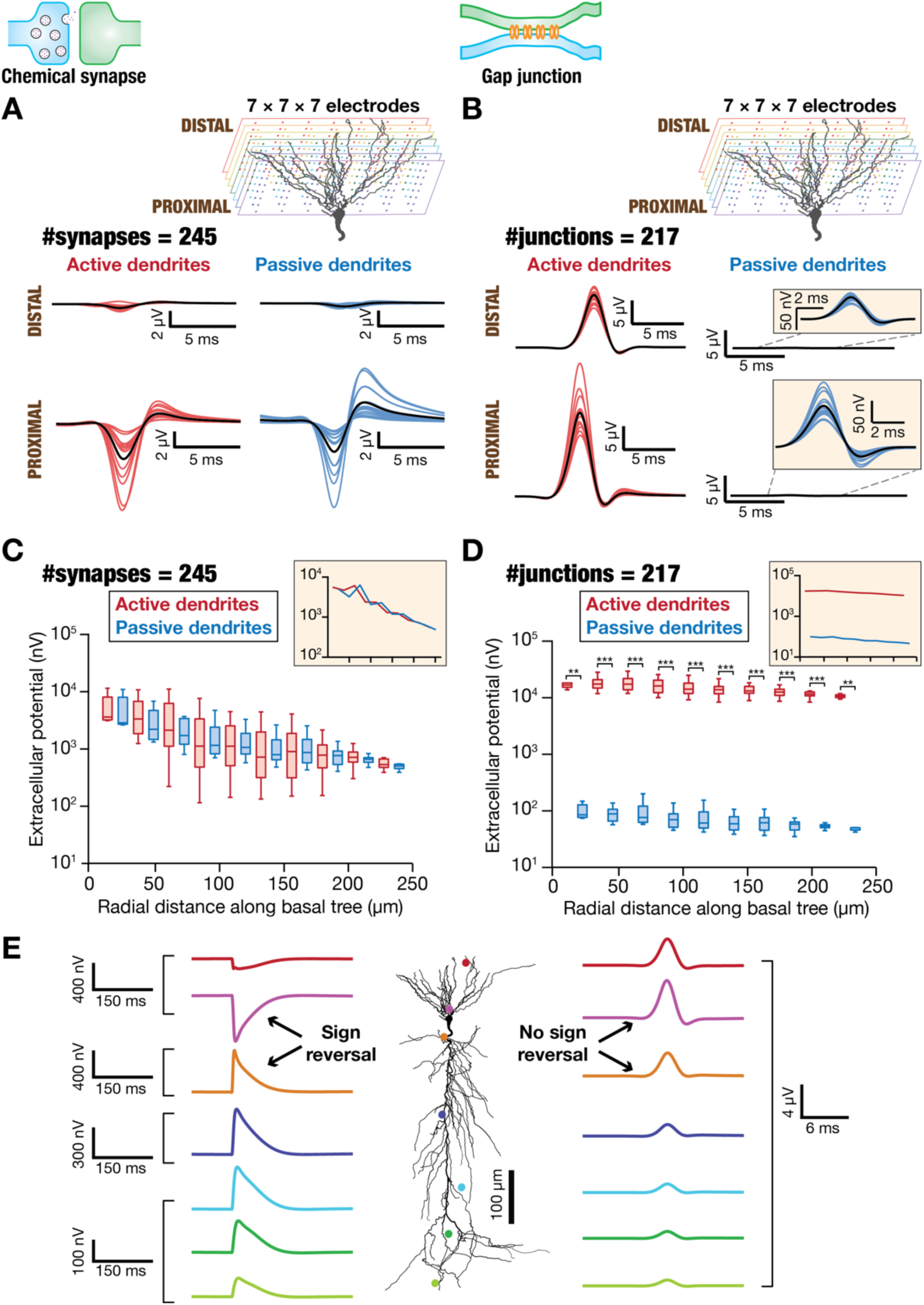
Contrasting extracellular signatures associated with active dendritic chemical synapses *vs*. gap junctions receiving synchronous inputs. *A. Top,* 3D electrode setup representing 7×7×7 (343 in total) electrode array spanning the basal dendrites of a CA1 pyramidal neuron morphology. Although the entire morphology was used for simulations spanning the apical and basal dendrites, the depiction here is restricted to the basal dendrites to emphasize electrode locations. *Bottom*, field potential traces from electrodes at proximal (15 traces representing different locations within 50–100 μm) and distal (21 traces representing different locations within 190–300 μm) locations along the somato-basal axis, when dendrites were active (*left*) or passive (*right*). Black traces depict the respective average trace across all distal or proximal locations. The 245 chemical synapses (*N*_syn_ = 245) which were randomly dispersed across the basal dendrites received synchronous inputs. *B.* Same as *A*, but for external inputs arriving through gap junctions. The number of gap junctions *N*_jun_ = 217. *C.* Amplitudes of negative deflection of field potentials for all 343 electrodes, plotted as functions of radial distance of the electrode from the soma, for active and passive dendritic models receiving synchronous inputs through chemical synapses (*N*_syn_ = 245). Inset shows plot of median field potential amplitude values as a function of distance for both active and passive dendritic configurations. *D.* Same as *C*, but amplitudes of positive deflections in extracellular potentials associated with inputs arriving through gap junctions. The number of gap junctions *N*_jun_ = 217. Comparison of active *vs*. passive dendritic configurations in (*C–D*): * *p*<0.05, \*\**p*<0.01, \*\*\**p*<0.001, Wilcoxon rank-sum test. *E.* Extracellular electrodes were placed across the entire span of the neuron (active dendrites with no sodium) instead of being confined to the basal dendritic span (panels *A–D*), with all parameters set identical to panels *A–D*. A flip in the sign of the extracellular potentials may be noted for synchronous stimulation with chemical synapses (*Left*), but not with stimulation with gap junctions (*Right*).

To assess the spatial profile of the extracellular potentials with synchronous stimulus configurations, we extended the placement of extracellular electrodes beyond the basal dendritic span to cover the apical dendrites of the morphology. When synchronous inputs arrived through chemical synapses, we found an expected shift in the sign of deflection in the extracellular potentials on the apical side of the dendritic morphology (which received no synaptic inputs) (Fig. 1*E*) indicative of a sink-source dipole. In contrast, when synchronous inputs arrived through gap junctions, the deflection in the extracellular potentials stayed positive across the entire stretch of the morphology (Fig. 1*E*). The magnitude of deflections showed expected reductions with increasing distance from the basal dendrites, irrespective whether inputs arrived onto basal dendrites through chemical synapses or gap junctions (Fig. 1*E*).

Together, these observations unveiled crucial distinctions in the polarity and the spatiotemporal patterns of extracellular potentials associated with synchronous inputs impinging on active dendrites through chemical synapses *vs*. gap junctions. Importantly, these results also pointed to the dominant role of contributions from active dendritic transmembrane currents in shaping extracellular field potentials associated with gap junctional inputs.

### Synchronous inputs: Outward transmembrane currents from active dendrites contribute to positive deflection in extracellular potentials associated with gap junctional inputs

Which of the several transmembrane currents contributed to the striking differences in the profiles of extracellular potentials associated with chemical synapses *vs*. gap junctions? In systematically addressing this question, we repeated our experiments with synchronous stimuli with models where specific currents were turned off. We assessed extracellular potentials in models without sodium and/or leak potassium currents and compared the outcomes with the default active dendritic model outcomes (Fig. 2). With synchronous inputs arriving through chemical synapses, these analyses revealed a role for sodium currents and action potential generation in determining the biphasic nature of the deflections in the extracellular potentials around basal dendritic regions (Fig. 2*A*). In the absence of sodium channels, extracellular potentials at all basal dendritic locations showed negative deflections, reflecting the kinetics of inward synaptic currents, with significantly reduced amplitudes (Fig. 2*A*). However, removal of leak currents did not yield pronounced changes in the extracellular potentials associated with synchronous inputs through chemical synapses (Fig. 2*A–B*). Along the basal dendritic neuropil, the values of peak negative deflection (Fig. 2*C*) and peak positive deflection (Fig. 2*D*) in the extracellular potentials reduced with electrode distance from the soma. The spatial profiles of deflections showed pronounced differences when sodium channels were removed but showed negligible differences with the removal of leak channels (Fig. 2*C–E*).

**Figure 2.**
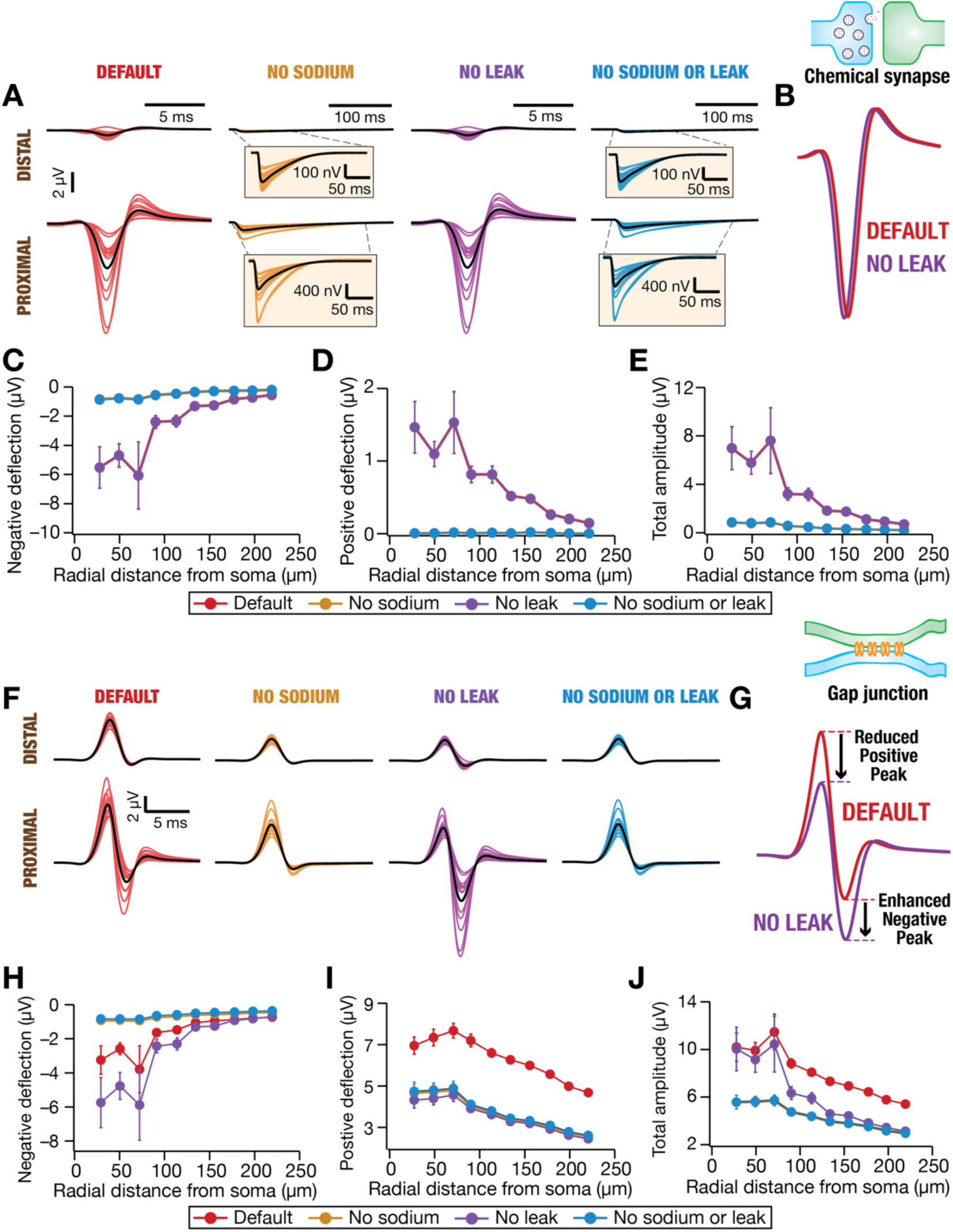
Differential polarity of field potentials associated with synchronous inputs through chemical synapses *vs*. dendro-dendritic gap junctions on active dendrites. *A*. Extracellular potentials from electrodes at proximal (15 colored traces represent different locations within 50–100 μm) and distal (21 colored traces represent different locations within 190–300 μm) locations along the somato-basal axis. Shown are traces for default (where all components were present), no sodium, no leak, and no sodium or leak scenarios for active dendritic structures. Black traces in each scenario depict the respective average trace across all distal or proximal locations. The 245 chemical synapses (*N*_syn_ = 245) which were randomly dispersed across the basal dendrites received synchronous inputs. *B*. Zoomed example trace (from a proximal electrode at 54 µm from soma) showing the impact of leak channels in shaping the extracellular potentials associated with active dendritic structures with (Default) and without leak channels (No leak)*. C–E.* Mean and SEM of the amplitudes of negative deflection *(C)*, positive deflection *(D)*, and the total peak to peak amplitude *(E)* of the extracellular potentials, plotted as functions of radial distance of electrode location, for default, no sodium, no leak, and no sodium or leak scenarios for active dendritic structures. *F–J*. Same as panels *A–E*, but for active dendritic structures receiving synchronous inputs through dendro-dendritic gap junctions (*N*_jun_ = 99).

In contrast, in models that received synchronous inputs through gap junctions, both sodium channels and leak channels played crucial roles in shaping the spatiotemporal profile of extracellular potentials (Fig. 2*F–J*). First, deleting sodium channels from the models prominently suppressed both negative and positive deflections in the extracellular potentials (Fig. 2*F*, Fig. 2*H–I*). Second, and in striking contrast with models activated by chemical synapses (Fig. 2*A–E*), deletion of leak channels enhanced the negative peak but reduced the positive peak of extracellular potentials when sodium channels were intact (Fig. 2*F–I*). Finally, when both sodium channels and leak channels were removed, the kinetics (Fig. 2*F*) and spatial profiles of deflections in extracellular potentials (Fig. 2*H–J*) were comparable to those where only sodium channels were removed.

These observations point to a critical role for a competition between the outward leak currents and the inward sodium currents in defining the spatiotemporal profile of extracellular potentials associated with synchronous inputs arriving through gap junctions. To elaborate, excitatory gap junctional inputs triggered depolarizing voltage responses in the postsynaptic neuron, which translated to outward currents through the leak and the other active channels. When this depolarization crossed action potential threshold, an action potential was generated by the confluence of inward sodium currents followed by outward currents through the potassium channels. Therefore, the outward transmembrane currents triggered by gap junctional inputs preceded the inward sodium current that generated the action potentials, resulting in a scenario where the positive peak in the extracellular potential preceded the negative peak (Fig. 2*F–G*).

When the sodium channels were deleted, the inward sodium current was suppressed and translated to a pronounced reductions in the negative (Fig. 2*H*) and the positive deflections (Fig. 2*I*) in the extracellular potentials (associated with gap junctional inputs). On the other hand, when leak channels were deleted, the outward leak current generated by gap-junctional depolarization and by the action potential were reduced (Fig. 2*F–G*). Together, such a scenario resulted in a reduced positive peak and an enhanced negative peak in the absence of leak channels but in the presence of sodium channels (Fig. 2*F–I*). Sodium channels were essential because they amplify gap junctional voltage depolarizations and mediate action potential generation. When sodium channels were absent, the outward currents triggered by gap-junctional depolarization dominated the extracellular potentials, with leak currents playing a minimal role in shaping extracellular potentials (compare “no sodium” *vs.* “no sodium or leak” cases in Fig. 2*F–J*).

What were the contributions of the different transmembrane currents to the extracellular potentials across the different scenarios presented above? We plotted the intracellular voltages and all transmembrane currents as functions of distance for each configuration assessed (Fig. 3). As expected, the presence of sodium channels yielded an action potential, irrespective of whether synchronous inputs came through chemical synapses (Fig. 3*A*) or through gap junctions (Fig. 3*C*). Specifically, the intracellular responses in the default and no leak models were comparable with synchronous inputs impinging onto active dendrites through either chemical synapses or gap junctions (Fig. 3 and Supplementary Fig. S3). In both models containing sodium channels (default and no leak), synchronous dendritic inputs reliably elicited action potentials regardless of the mode of synaptic transmission exhibiting similar electrical responses. The presence of an action potential translated to large driving forces for all ionic currents, including currents through synaptic receptors (Fig. 3*A–B*, Fig. 3*D*). In addition, the large temporal derivate associated with action potentials also translated to higher values of capacitive current in the presence of sodium channels (Fig. 3*B*, Fig. 3*D*). The higher leak currents in the presence of sodium channels (Fig. 3*B*, Fig. 3*D*) contributed to the shifts in negative and positive peaks in extracellular potentials that were triggered by deletion of leak channels (Fig. 2*F–J*).

**Figure 3.**
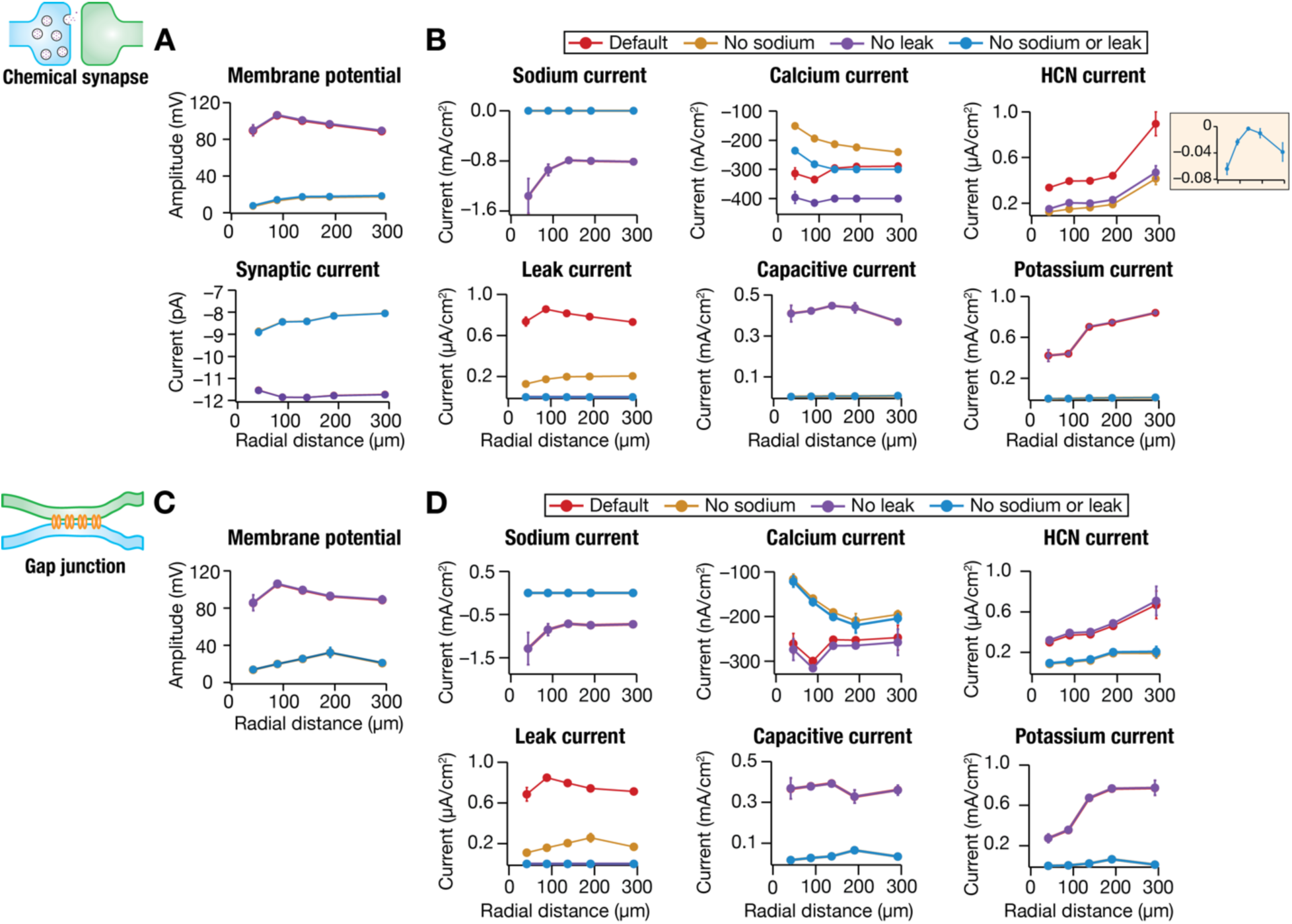
Ionic basis of the differential contributions of transmembrane currents to field potentials associated with synchronous inputs arriving through chemical synapses *vs*. gap junctions on active dendrites. *A*. Mean and SEM of peak membrane voltages (*top*) and peak synaptic currents (*bottom*) from across somato-basal locations recorded intracellularly for all 4 model configurations. The 4 different model configurations shown are the default active model, and the active models where sodium channels, leak channels, or both sodium and leak channels were absent. It may be noted that there were no action potentials or dendritic spikes when there were no sodium channels in the models. The dependence of synaptic current on the membrane potential, acting as the driving force, may also be noted. *B*. Mean and SEM of peak values of transmembrane sodium, calcium (*T*-type, *L*-type, *R*-type, and *N*-type), HCN, leak, capacitive, and potassium (*A*-type, delayed rectifier, and *M*-type) currents for different active models receiving synchronous inputs through chemical synapses, plotted as functions of radial distance from soma for all 4 model configurations. *C–D*. Same as panels *A–B*, but with different configurations of active models receiving synchronous inputs through gap junctions. There are no synaptic currents plotted here as there are no transmembrane synaptic currents associated with gap junctions.

An important observation here is related to the current through HCN channels in the presence of abrupt large-amplitude voltage deflections, such as those observed with synchronous synaptic input. We noted that except for one case (“No sodium or leak” with chemical synapses), all average HCN current values were positive in sign (Fig. 3*B*, Fig. 3*D*). How do HCN channels, which typically carry inward transmembrane currents that are negative in sign, yield positive outward currents? HCN channels are active at rest and are endowed with slow activation/deactivation kinetics with a time constant ranging from tens to hundreds of milliseconds (Robinson and Siegelbaum, 2003; Biel et al., 2009; Shah, 2014; Combe and Gasparini, 2021; Mishra and Narayanan, 2025). With synchronous inputs impinging on the neuron, the transition from rest to large depolarization (which trigger action potentials in the presence of sodium channels) is fast. The fast kinetics of the depolarization implies that the active fraction of slow HCN channels does not have sufficient time for deactivation. Thus, once the membrane voltage crosses the reversal potential of HCN channels (*E_h_* = – 30 mV), HCN current (*I_h_* = *g*_HCN_(*V* − *E_h_*)) becomes positive as the driving force (*V* − *E_h_*) becomes positive and the resting conductance remains non-zero. Together, in scenarios where the depolarizing voltage kinetics is faster than the deactivation kinetics of HCN channels towards crossing *E_h_*, HCN channels yield an outward current (Supplementary Fig. S4). It is important to note that despite their relatively small magnitude, the outward HCN currents (Fig. 3*D*) substantially contribute to positive extracellular potential deflections associated with gap junctional inputs (Fig. 2), together with leak and other outward currents.

While outward HCN currents (Fig. 3*B*) are also expected to influence LFPs under chemical synaptic input, their impact was minimal due to the predominance of large inward synaptic currents (Fig. 3*A*). As LFPs reflect the summation of all transmembrane currents, the dominant contributors vary across different modes of synaptic transmission.

Together, our results unveil stark differences in the extracellular field potentials associated with synchronous inputs, depending on whether such inputs arrived onto active dendrites through chemical synapses or gap junctions. Whereas leak channels played minimal roles in shaping extracellular potentials when inputs arrived through chemical synapses, they played a prominent role in defining extracellular potentials when inputs arrived through gap junctions. Our results showed that the striking polarity differences observed in extracellular potentials associated with chemical synapses *vs*. gap junctions (Fig. 1*A–B*) were driven by two prominent factors: (i) the absence of inward transmembrane currents with gap junctional inputs; and (ii) the presence of outward currents (through leak, potassium, and HCN channels) associated with gap-junctional depolarization and action potentials triggered by such depolarization (Figs. 2–3).

### Random inputs: Dichotomy in extracellular potentials associated with asynchronous inputs through chemical synapses *vs*. gap junctions

We next explored if there were distinctions between extracellular potentials stemming from random inputs through chemical *vs.* electrical synapses. We computed extracellular potentials from models receiving low– (Fig. 4) and high-frequency (Supplementary Fig. S5) random inputs (LFRI and HFRI) onto active dendrites through chemical synapses or gap junctions. For both LFRI and HFRI, the power associated with extracellular potentials was higher when asynchronous inputs arrived through chemical synapses compared to their arrival through gap junctions (Fig. 4; Supplementary Fig. S5). These observations held irrespective of whether models were endowed with sodium channels or not (Fig. 4; Supplementary Fig. S5). Across scenarios, the power associated with extracellular potentials reduced with increasing distance from the cell body (Fig. 4; Supplementary Fig. S5). We attributed these differences to the presence of an additional transmembrane current through the synaptic receptors, in models that received inputs through chemical synapses when compared to models receiving inputs through gap junctions. Importantly, the presence of the large inward synaptic currents in structures receiving chemical synaptic inputs also translated to predominantly negative deflections in associated extracellular potentials. In comparison, extracellular potentials associated with models that received gap junctional inputs showed deflections on both positive and negative directions. This distinction in the sign of the extracellular potentials associated with chemical synapses *vs*. gap junctions was prominently visible in models where there were no sodium channels (Fig. 4; Supplementary Fig. S5). Together, these results revealed distinctions in extracellular potentials associated with active dendritic structures receiving random inputs at different frequencies through chemical synapses *vs*. gap junctions.

**Figure 4.**
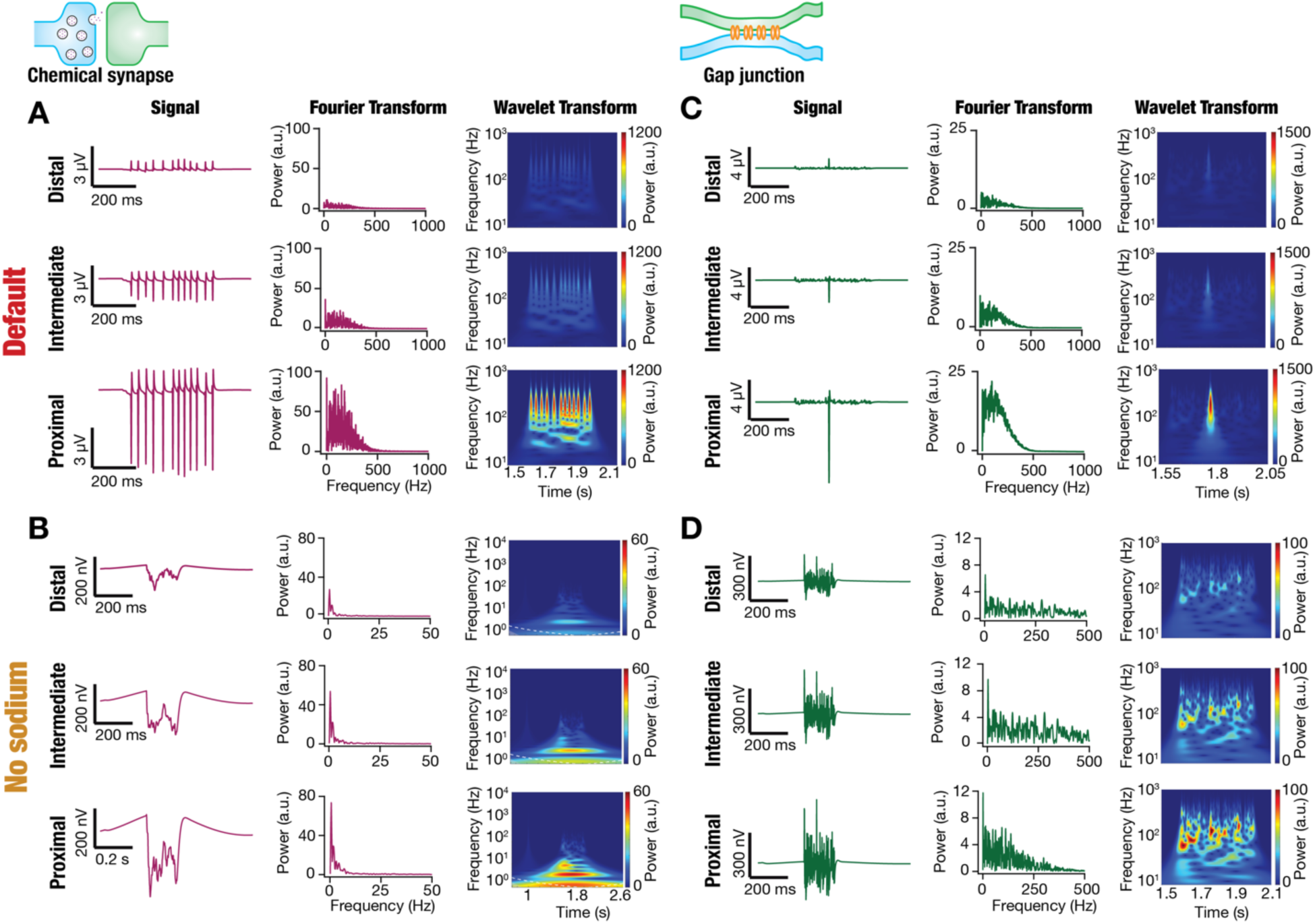
Differential spatiotemporal structure of field potentials associated with active dendrites receiving low-frequency random inputs through chemical synapses *vs.* gap junctions. *A*. Distance-wise LFP responses to low-frequency random inputs (LFRI) impinging on active dendrites through chemical synapses. *Rows 1–3:* LFP data from electrodes located at a distal (∼152 µm; *Row 1*), intermediate (∼97 µm; *Row 2*), and proximal (∼55 µm) locations with reference to their radial distance from the soma. *Column 1:* time-domain signal. *Column 2:* Fourier transform of the signal shown in *Column 1*. *Column 3:* spectrogram of the signal shown in *Column 1* computed using wavelet transform. *B*. Same as panel (*A*) but for active model lacking sodium conductance. *C–D*. Same as panels (*A–B*), except low-frequency random inputs impinged onto active dendrites through gap junctions.

### Rhythmic oscillatory inputs: Distinctions in spectral characteristics of extracellular potentials associated with chemical synapses *vs*. gap junctions on active dendrites

In a third set of experiments, we activated the chemical synapses or the gaps junctions with rhythmic inputs at different physiologically relevant frequencies (Fig. 5). We computed the associated extracellular potentials and studied their spectral properties in models that were endowed with active or passive dendrites (Fig. 5). As expected, the extracellular potentials manifested oscillatory patterns in the same range as that of the stimulation frequencies, irrespective of whether the inputs came through chemical synapses or gap junctions and irrespective of whether the dendrites were active or passive. We computed the peak power from the Fourier spectra of extracellular potentials from all 343 electrodes for each scenario, involving different frequencies (spanning 1–128 Hz in octaves), model configurations (active *vs*. passive dendrites), and modes of stimulation (through chemical synapses *vs*. gap junctions) (Fig. 6).

**Figure 5.**
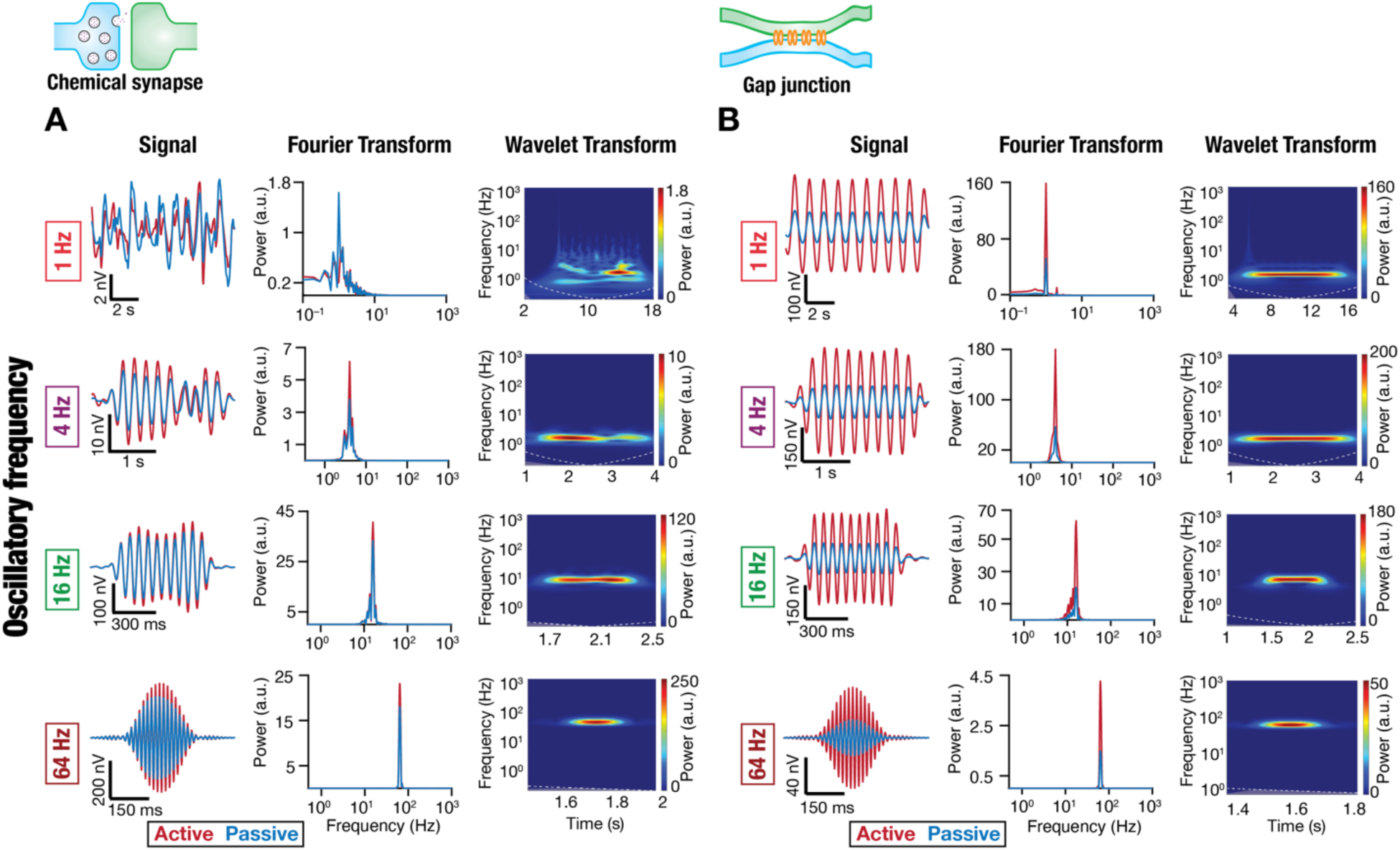
Dominance of specific oscillatory bands in field potentials depended on whether inputs onto active dendrites were received through chemical synapses or gap junctions. *A*. Example LFP responses to rhythmic inputs at different (1–64 Hz) frequencies and their spectral signatures, shown for simulations performed with active or passive basal dendrites receiving rhythmic inputs through chemical synapses. Each row shows the filtered LFP signal at an electrode placed ∼97 µm from the soma, the Fourier power spectrum, and the wavelet spectrogram for the LFP signal. Different rows depict different input frequency values for the rhythmic input (1 Hz, 4 Hz, 16 Hz, and 64 Hz). *B*. Same as *(A)* but for the rhythmic inputs impinging on basal dendrites at different frequencies through gap junctions. All simulations depicted here were performed in the absence of sodium channels to avoid spiking.

**Figure 6.**
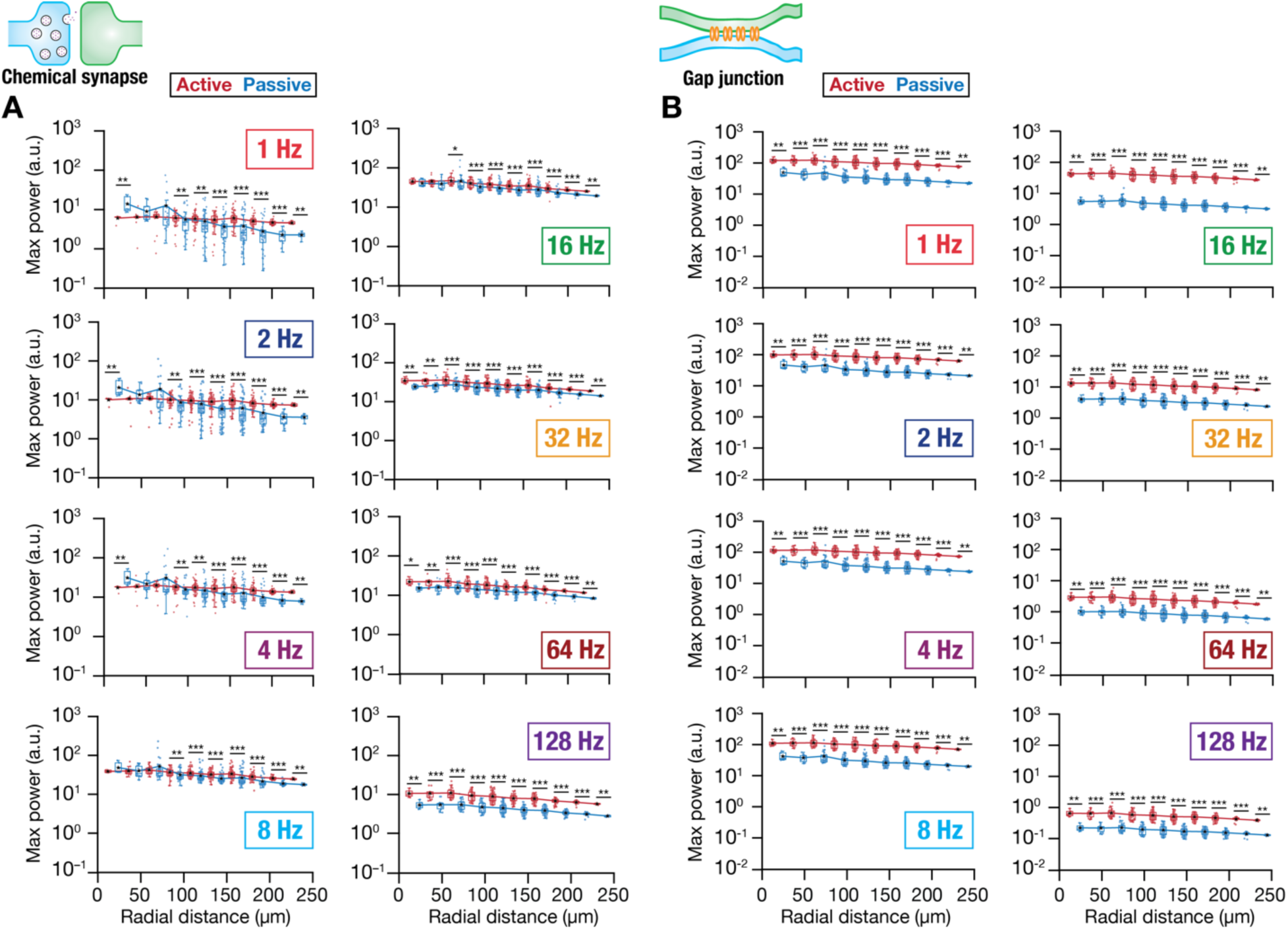
Distance-dependence of spectral power in specific bands of field potentials associated with active dendrites receiving rhythmic inputs through chemical synapses or gap junctions. *A*. Maximum power in LFP responses associated with rhythmic inputs at different (1–128 Hz) frequencies, shown for simulations performed with active or passive basal dendrites receiving these rhythmic inputs through chemical synapses. All electrodes at specific radial distances are depicted for each scenario. The frequency of the rhythmic input is highlighted in each panel. *B*. Same as *(A)* but for the rhythmic inputs impinging on basal dendrites at different frequencies through gap junctions. Across all plots, lines connect the respective median values (represented by black stars). All simulations depicted here were performed in the absence of sodium channels to avoid spiking. * *p*<0.05, \*\**p*<0.01, \*\*\**p*<0.001 (Wilcoxon rank-sum test)

As with other stimulus configurations before, the presence of an additional transmembrane current with chemical synaptic stimulation contributed to differences in extracellular potentials associated with chemical synapses and gap junctions. First, these differences implied that there were dominant contributions from active dendritic transmembrane currents to extracellular field potentials associated with gap junctional inputs (Fig. 6*B*), but not with chemical synaptic inputs (Fig. 6*A*). This was inferred from the pronounced differences observed in active *vs*. passive model configurations with gap junction inputs (Fig. 6*B*), but not with chemical synaptic inputs (Fig. 6*A*; except for 128 Hz where pronounced differences were noted for both chemical synapses and gap junctions). Specifically, across frequencies, the LFP power with active dendrites was larger than LFP power with passive dendrites when inputs impinged through gap junctions (Fig. 6*B*), implying a significant contribution from active dendritic transmembrane currents.

Second, the dominant role of the receptor currents with chemical synaptic stimulation implied that there was a strong transmembrane component spanning all frequencies. Thus, we observed relatively higher power even at higher frequencies (say, for 64 Hz or 128 Hz) in the extracellular potentials (as well as total transmembrane currents) with chemical synaptic stimulation (Fig. 7*A–B*) compared to those with gap junctional stimulation (Fig. 7*C–D*). With gap junctional rhythmic inputs, the power of extracellular potentials (and total transmembrane currents) at higher frequencies was low (Fig. 7*C–D*), owing to their origins being entirely through transmembrane currents driven by a membrane-filtered voltage waveform. Specifically, as the membrane resistor-capacitor circuit acts to suppress higher frequencies, intracellular voltage responses to high-frequency current inputs were lower in amplitude yielding low-amplitude transmembrane currents due to low driving forces. Although such filtering is common to both chemical synaptic and gap junctional inputs, the reliance of LFP associated with gap junctional inputs on voltage-driven transmembrane currents translate to the distinctions observed in extracellular signatures associated with chemical synapses (Fig. 7*A–B*) *vs*. gap junctions (Fig. 7*C–D*).

**Figure 7.**
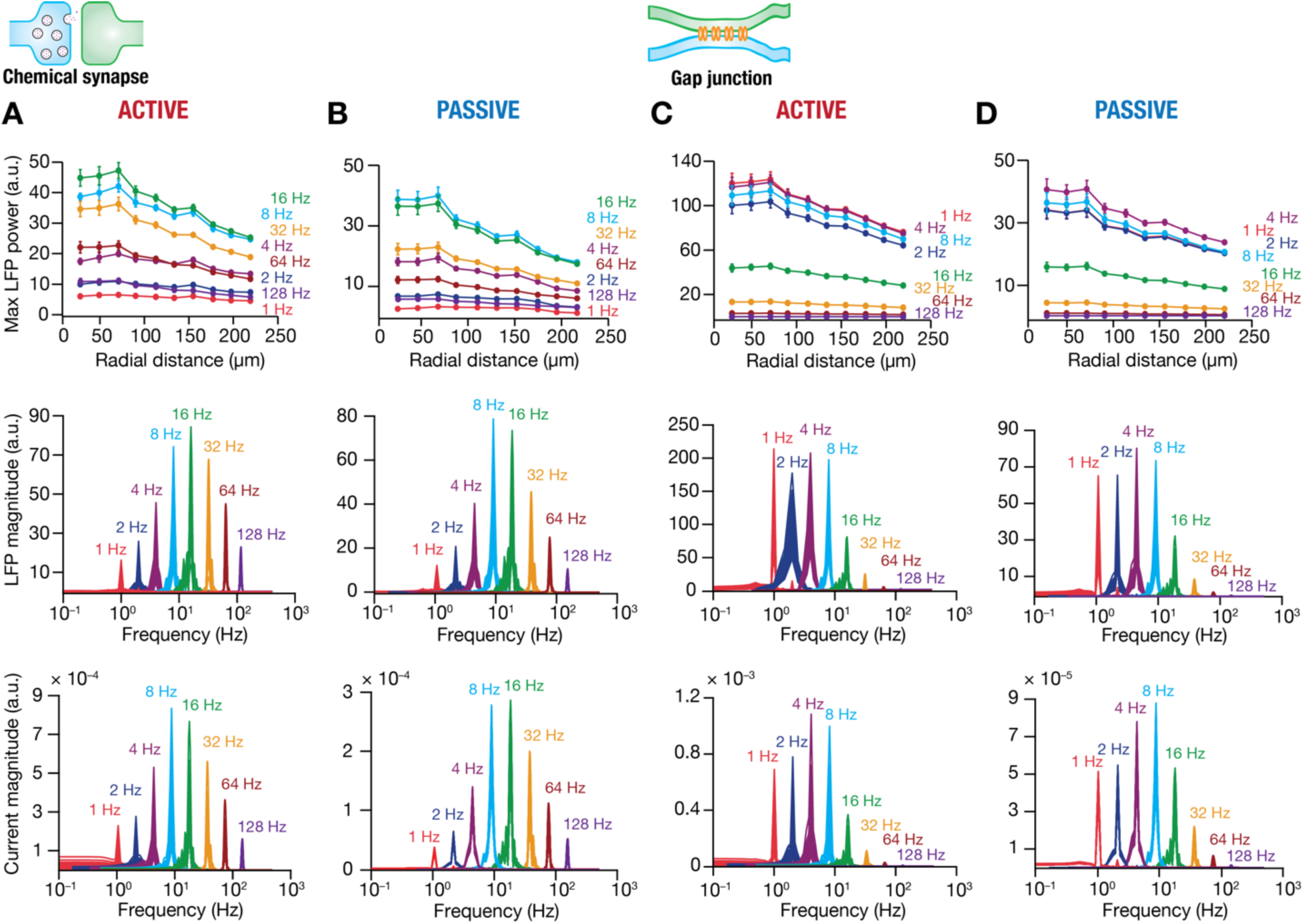
Transmembrane currents driven by voltage responses mediate the differential emphasis of specific oscillatory bands in field potentials associated with chemical synapses *vs*. gap junctions on active dendrites. *A. Row 1:* Distance-dependent maximal power of local field potentials recorded at different electrodes (shown as mean and SEM) associated with neuronal response to rhythmic inputs at different frequencies impinging on the active basal dendritic model through chemical synapses. *Row 2:* Fourier power spectra for all field potentials at different frequencies of the rhythmic inputs. Each trace for a given frequency represents different electrodes. *Row 3:* Fourier spectra of the filtered total transmembrane current for each frequency of rhythmic inputs, from each basal dendritic compartment. *B.* Same as panel *A*, but for simulations performed with passive dendrites. *C–D.* Same as *(A–B)*, but with rhythmic inputs coming through gap junctions. All simulations depicted here were performed in the absence of sodium channels to avoid spiking.

Third, the differences in the contributions of active transmembrane currents *vs*. receptor currents to the LFP also translated to a differential emphasis of frequencies associated with gap junctions *vs*. chemical synaptic stimulations (Fig. 7*A–D*). Specifically, rhythmic inputs at 8 Hz and 16 Hz showed maximal power in the extracellular potentials associated with chemical synapses, irrespective of whether the dendrites were active or passive (Fig. 7*A–B*). On the other hand, with gap junctional rhythmic stimulation, we found that extracellular potentials associated with lower frequencies (1–8 Hz) manifested much higher power compared to the higher frequencies, again irrespective of whether the dendrites were active or passive (Fig. 7*C–D*). Importantly, there was minimal reduction in power of extracellular potentials across most frequencies (except at 128 Hz) between active (Fig. 6*A*, Fig. 7*A*) and passive (Fig. 6*A*, Fig. 7*B*) dendritic configurations with chemical synaptic stimulation. In striking contrast, there was a large reduction in power when active components were removed from the dendrites as rhythmic inputs arrived through gap junctions (Fig. 6*B*, Fig. 7*C* vs. Fig. 7*D*). These observations reaffirm the relative dominance of receptor currents and active transmembrane currents for chemical synaptic and gap junctional stimulus conditions, respectively.

Fourth, we looked at phase differences in extracellular potentials across the neuronal morphology to span apical dendrites as well (Supplementary Fig. S6). We found important differences in the spatiotemporal profiles of extracellular potentials obtained with rhythmic inputs arriving through chemical synapses *vs*. gap junctions. With active dendritic structures, we found a progressive shift in the phase of extracellular potentials as the electrode location moved away from the basal dendrites where the chemical synaptic inputs were impinging (Supplementary Fig. S6*A*). However, there was negligible phase difference across electrode locations when inputs onto basal dendrites arrived through gap junctions (Supplementary Fig. S6*A*). With passive dendritic structures, a similar relative shift in phase was observed between chemical synaptic *vs*. gap junctional inputs. However, a smaller phase shift was observed across electrodes when oscillatory inputs arrived through gap junctions (Supplementary Fig. S6*B*).

Together, we observed several crucial distinctions in the spectral characteristics of extracellular potentials associated with gap junctional *vs*. chemical synaptic rhythmic inputs impinging on active dendrites. We found these distinctions to be attributable to the relative contributions of receptor *vs*. active transmembrane currents to extracellular potentials associated with chemical synaptic *vs*. gap junctional inputs.

### Rhythmic oscillatory inputs: LFP-spike phase relationship relied on the oscillatory frequency and on whether inputs arrived through chemical synapses or gap junctions

The phase of spikes in individual neurons with reference to extracellular potentials are studied in phase coding schemas and in assessing synchrony. Given key differences in the extracellular spectral signatures (Figs. 5–7) and spatiotemporal profiles of LFPs (Supplementary Fig. S6), we asked if there were differences in the phase relationship between spikes and extracellular potentials when rhythmic inputs arrived through chemical synapses *vs*. gap junctions (Fig. 8; Supplementary Figs. S7–S12). We used extracellular potentials from a soma-proximal electrode as reference and computed phase of somatic action potentials, considering trough of the LFP trace as zero degrees. These experiments and analyses were repeated across different trials (which were different in terms of synaptic localization profiles), each spanning multiple cycles, with rhythmic inputs arriving at different frequencies through chemical synapses or gap junctions onto different model configurations (Fig. 8; Supplementary Figs. S7–S12). Across frequencies and irrespective of whether inputs arrived through chemical synapses or gap junctions, spikes manifested phase locking to LFP oscillations (Fig. 8*A–B*; Supplementary Figs. S7–S8). There were frequency-dependent differences in spike phases, depending on whether rhythmic inputs arrived through chemical synapses or gap junctions (Fig. 8*A–B*; Supplementary Figs. S7–S8). Irrespective of whether inputs arrived through chemical synapses or gap junctions, the median spike phase (across trials and cycles) progressively shifted from ∼180° to ∼0° with increase in frequency from 1 Hz to 128 Hz (Fig. 8*B*). However, for intermediate frequency values (8 Hz, 16 Hz), there were large differences in the spike phase values for models receiving inputs through chemical synapses *vs*. gap junctions.

**Figure 8.**
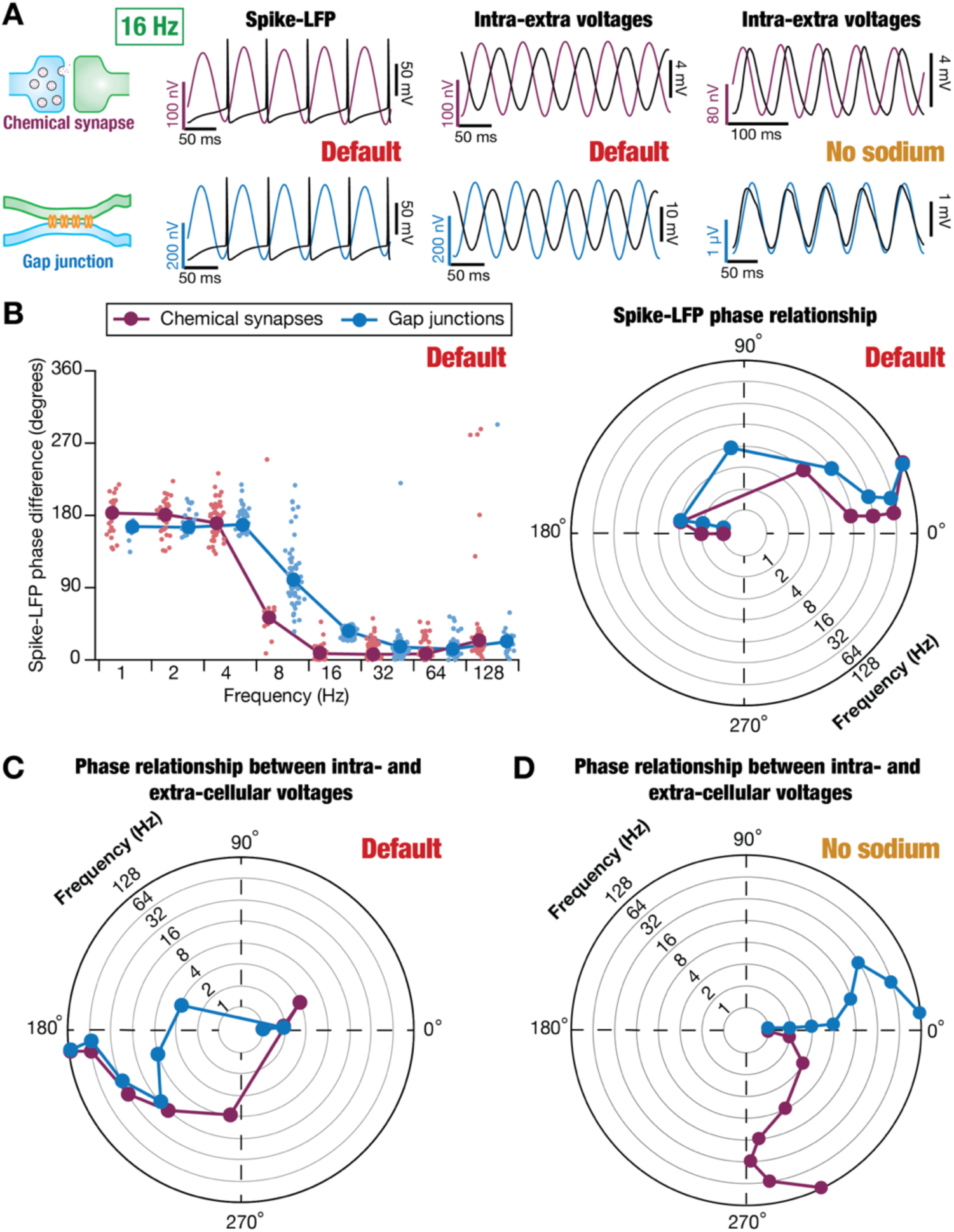
Differential phase relationship between local field potentials and spikes associated with rhythmic inputs through chemical synapses *vs.* gap junctions on active dendrites. *A.* Example LFP traces (color coded based on whether they are associated with chemical synapses in dark pink or gap junctions in blue) and simultaneously recorded intracellular somatic voltage traces (Black) for neurons receiving rhythmic inputs through chemical synapses (*top*) or gap junctions (*bottom*). Shown are traces with default model configuration where all channels were intact (*left*), traces where the intracellular traces were filtered to the respective band (*middle*), and extracellular/intracellular traces obtained in the absence of sodium channels (*right*). *B. Left,* Spike phase with reference to local field potentials for each spike (lighter circles) for oscillatory inputs at different frequencies impinging on active dendrites through chemical synapses *vs.* gap junctions. Dark-colored circles represent the median values at each frequency for the respective group*. Right,* Polar version of the plot showing median of spike-LFP phases over five trials for rhythmic inputs at different frequencies through chemical synapses and gap junctions. The different frequencies are represented along the concentric circles and the corresponding spike-LFP phase values are plotted along the angular axis. *C.* Polar plot with median of phases obtained from cross-correlation of *filtered* intracellular potential and corresponding LFP traces at respective frequencies over all trials. These plots were derived from the same intracellular traces as in panel *A* but represent phase differences between extracellular traces and the entire filtered intracellular voltage trace. *D.* Polar plot of phases obtained from cross-correlation between intracellular potential (without sodium conductance) and corresponding LFP traces when oscillatory inputs were presented through chemical synapses or gap junctions.

To understand these spike phase profiles across frequencies and across type of synapses, we compared the phase difference between intracellular somatic voltage waveforms and peri-somatic extracellular potentials. We filtered the waveforms with reference to the specific frequency of rhythmic inputs they were obtained in response with (Supplementary Table S3) and performed cross-correlation analysis to compute phase differences between intra– and extra-cellular potentials (Fig. 8*C*; Supplementary Figs. S9–S10). We repeated these analyses across different trials and phase differences were plotted. Consistent with what we observed with spike phases earlier, we found that the phase difference between extra– and intra-cellular potentials progressively shifted from ∼0° to ∼180° as frequency increases. Note that a spike that occurs at the peak of the extracellular potential (with the trough designated as 0°) would be assigned a spike phase of 180°. Under this scenario, the peak of intracellular voltage would match with the peak of extracellular waveform, thereby translating to a 0° phase difference between intra– and extracellular voltages. We observed negligible differences in this phase relationship for chemical synapses *vs*. gap junctions for low (1–2 Hz) and high frequencies (16–128 Hz). However, for intermediate frequency values (4–8 Hz), there were differences in the phase relationship between intra– and extra-cellular potentials depending on whether rhythmic inputs arrived through chemical synapses or gap junctions (Fig. 8*C*; Supplementary Figs. S9–S10).

Finally, to assess the specific contributions of the spike generation and sodium channels to these phase relationships, we repeated our cross-correlation analyses on simulations that were performed in the absence of sodium channels (Fig. 8*D*; Supplementary Figs. S11–S12). In this scenario, whereas the phase relationships between intra– and extra-cellular voltages were comparable for low frequencies (1–2 Hz), there was a dramatic deviation between phase plots obtained with chemical synaptic *vs*. gap junctional inputs (Fig. 8*D*; Supplementary Figs. S11–S12). Specifically, at higher frequencies (observed maximally at 16 Hz), the intracellular voltage waveform lagged the extracellular voltage with chemical synaptic rhythmic stimulation (Fig. 8*D*; Supplementary Figs. S11). In contrast, the intracellular waveform manifested a small lead with reference to the extracellular waveform with gap junctional rhythmic stimulation (Fig. 8*D*; Supplementary Figs. S12). In either case, the magnitude of lead or the lag never crossed 90° for any of the analyzed frequencies.

These observations point to a scenario where the depolarization induced by the receptor currents trigger the activation of outward currents, which dominate the extracellular potentials with chemical synaptic stimulation in the absence of sodium channels (Fig. 8*D*; Supplementary Fig. S11). The transmembrane inward receptor current and the outward currents together resulted in a phase lead in the LFP observed with chemical synaptic stimulation, especially with higher frequencies (Fig. 8*D*; Supplementary Figs. S11). On the other hand, with gap junctional stimulation, depolarization triggered by gap junctional currents result in net outward currents following the intracellular depolarization caused by the rhythmic inputs. Such a scenario translates to a lag in the extracellular voltage trace compared to its intracellular counterpart (Fig. 8*D*; Supplementary Fig. S12). In the presence of sodium channels, the large voltage deflections and spikes generated, the consequent increase in driving force, the inward transmembrane sodium current, the activation-inactivation profiles of all active channels, the spatiotemporal profile of stimulation, the membrane time constant, and the specific kinetics of the synaptic activation together result in a scenario where spike phases are variable for gap junctions *vs*. synaptic stimulation (for intermediate frequencies).

## DISCUSSION

A diverse interplay involving several factors — such as neuronal morphology, organization of different morphologies and afferent inputs onto them, active components in different neuronal processes, spatiotemporal patterns of afferent inputs onto neurons, receptor localization and kinetics, intraneuronal filtering, neuropil properties, ephaptic coupling, interactions among cross-cellular transmembrane currents, and electrode localization — shapes extracellular potentials. Our analyses add a further layer of complexity by demonstrating stark differences in extracellular potentials associated with inputs arriving onto active dendrites through chemical synapses *vs*. gap junctions. Considering the crucial role of gap junctions in several networks across the brain, it is critical that LFP analyses and interpretations account for whether inputs arrive through gap junctions or through chemical synapses. Our analyses place strong emphasis on accounting for the distinct contributions of gap junctional inputs onto active structures (*e.g.*, neuronal dendritic and axonal compartments, neuronal and glial soma, glial processes) to extracellular field potentials. The absence of receptor transmembrane currents and the dominance of voltage-driven passive/active transmembrane currents in determining extracellular potentials associated with gap junctional inputs are crucial factors that must be accounted for in interpreting extracellular potentials. Our study emphasizes the need for quantitative assessment of how chemical and electrical synapses individually influence extracellular patterns across different brain regions and how their combined impact on extracellular potentials contribute to specific aspects of network physiology and accurate interpretations of brain signals.

### Dominance of active dendritic currents with LFP associated with gap junctions

When dendrites were thought to be passive, LFPs were considered to be predominantly reflective of currents through synaptic receptors that were present on different postsynaptic locations on neurons. However, it is now clear that dendrites are active in that they are endowed with several voltage– and/or calcium-gated ion channels. These dendritic active currents make predominant contributions to LFP apart from receptor currents. The receptor current, all active currents, the capacitive and leak currents across the neuronal morphology obey Kirchoff’s current law (KCL), thus creating source-sink dipoles depending on the open or closed nature of the field organization (Buzsaki et al., 2012; Einevoll et al., 2013; Reimann et al., 2013; Sinha and Narayanan, 2015; Ness et al., 2016; Sinha and Narayanan, 2022; Halnes et al., 2024).

However, much of these analyses have been limited to scenarios where synaptic inputs are received exclusively through chemical synapses, which manifest a transmembrane current through the postsynaptic receptors. With electrical synapses, the KCL is balanced without a transmembrane synaptic current because gap junctional inputs are direct intracellular inputs into the neuron that do not manifest a direct transmembrane current. Therefore, with chemical synapses, the synaptic current provides an extracellular field component that precedes the voltage response to the synaptic current. The voltage deflection caused by chemical synaptic or gap junctional inputs would yield additional transmembrane currents in the form of the leak current, the capacitive current, and the active currents. These additional transmembrane currents from different neuronal locations summate at an electrode location weighted in distance-dependent manner to eventually yield the extracellular potential. While these voltage-response-induced currents are common to both chemical synapses and gap junctions, the absence of transmembrane synaptic currents with gap junctional inputs results in important and striking differences in the extracellular signatures associated with gap junctions and chemical synapses. If dendrites were passive, the only transmembrane current components associated with gap junctional inputs would have been the leak and the capacitive currents. But, in a scenario where dendrites are active, the extracellular field was defined by active transmembrane currents as well, together contributing to the notable distinctions reported here.

In the absence of transmembrane synaptic currents, the outward currents that were triggered by depolarization dominated the extracellular field potentials with synchronous synaptic inputs. There was a striking difference in polarity of the field potentials associated with chemical synapses and gap junctions. These differences resulted a scenario where there was no source-sink dipoles in extracellular potentials across the neuronal morphology because gap junctions imply that inputs were injected directly into the neuron. The shape of the extracellular potentials, their polarities, and their spatial distributions have all been typically understood from the perspective of synaptic potentials. Our study demonstrates that these analyses do not hold for active structures connected through gap junctions.

Our analyses quantitatively demonstrates that the fundamental difference between the gap junctional and chemical synaptic contributions to extracellular potentials is the absence of the direct transmembrane component from gap junctional inputs. A multitude of factors suggests that the observed LFP differences result not from variations in input strength or patterns but rather from differences in direct transmembrane currents, which are subsequently reflected in the LFP signals. *First*, the inputs were distributed randomly across the basal dendrites, irrespective of whether they were through gap junctions or chemical synapses. For both chemical synapses and gap junctions, the inputs were exclusively excitatory in nature.

*Second*, the results remained consistent regardless of the number of gap junctions or chemical synapses. (Fig. 1 with 217 gap junctions or 245 chemical synapses and Supplementary Fig. 2 with 99 gap junctions or 30 chemical synapses). Our fundamentally novel result that gap junctions onto active dendrites shape LFPs holds true irrespective of the relative density of gap junctions onto the neuron.

*Third*, for synchronous chemical synaptic inputs, the waveforms resembled typical postsynaptic potentials. Whereas, for gap junctional inputs, the waveforms showed characteristics of postsynaptic potentials or dendritic spikes (accounting the active nature of inputs from the potential presynaptic cells). Electrical response of postsynaptic cell remains identical, producing an action potential regardless of whether inputs arrive via gap junctions or chemical synapses. We quantitatively matched the strengths such that the model generated a single action potential in response to synchronous inputs, irrespective of whether they arrived through chemical synaptic or gap junctional inputs. We mechanistically analyzed the contributions of different cellular components and show that the direct transmembrane current in chemical synapses is the distinguishing factor that determines the dichotomy between the contributions of gap junctions *vs.* chemical synapses to extracellular potentials (Fig. 2-3).

*Fourth*, for random inputs, the models were not specifically tuned to generate a single action potential. Here, the inputs served as a proxy for asynchronous inputs arriving from other subregions at random times. *Finally*, the intracellular responses were comparable for chemical synaptic and gap junctional rhythmic inputs (Supplementary Fig. S6–S12). Here, the model was tuned to elicit a single spike per cycle in simulations evaluating spike-LFP interactions, ensuring a fair comparison between LFPs from gap junctional and chemical synaptic inputs.

Our results emphasize the need to revisit the frameworks for understanding field potentials, especially in brain regions where there are strong gap junctional connectivity across neurons (Traub, 1995; Bennett and Zukin, 2004; Connors and Long, 2004; Sohl et al., 2005; Traub et al., 2018), including scenarios where there is mixed connectivity involving gap junctions as well as chemical synapses (Hamzei-Sichani et al., 2012; Nagy et al., 2019).

### High-frequency LFP power was suppressed with gap junctional inputs

The absence of transmembrane receptor currents with gap junctional stimulation implies that membrane filtering has a dominant impact on extracellular potentials when synaptic inputs arrive through gap junctions. A direct consequence of such strong dependence on membrane filtering, which predominantly acts to suppress high frequencies, is that there is pronounced attenuation of extracellular potentials associated with high-frequency rhythmic inputs arriving through gap junctions (Figs. 5–7). Although high-frequency chemical synaptic inputs also face the same membrane filtering, the transmembrane receptor component provides an important additional contribution to extracellular potentials beyond the voltage-driven transmembrane currents.

There are several implications to this. First, although there might be strong high-frequency interactions between neurons coupled through gap junctions, the extracellular signatures associated with such strong interactions would be weak unless these interactions produce action potentials and translate to large extracellular signatures. Second, oscillatory networks in low frequency (say, delta or theta frequency ranges) connected through gap junctions might manifest strong extracellular signatures associated with rhythmic inputs. The strong dependence on membrane filtering implies that the extracellular potentials might not be of high amplitude if the oscillatory inputs through gap junctions were in the high-frequency ranges (say high-gamma or ripple band oscillations). In this context, our analyses lay the foundation for network analyses of the impact of gap junctions on LFPs. The conclusions from the single-neuron simulations in this study extend to a neuropil with several neurons, each receiving synaptic and gap junctional inputs, especially with reference to high-frequency ripple oscillations (Sirmaur and Narayanan, 2024). A neuropil made of hundreds of morphologically realistic pyramidal neurons was used to assess the role of different synaptic inputs — excitatory, inhibitory, and gap junctional — with different patterns of stimulation and active dendritic contributions to ripple-frequency oscillations. Network-based analyses have unveiled a dominant mediatory role of patterned inhibition in ripple generation, with recurrent excitations through chemical synapses and gap junctions, in conjunction with return-current contributions from active dendrites, playing modulatory roles in governing ripple characteristics (Sirmaur and Narayanan, 2024). Future studies could expand on these conclusions to explore the implications of frequency-dependent filtering (with reference to gap junctional coupling) on high-frequency extracellular oscillations.

Third, the extent of spatial spread of field potentials, especially in low frequencies, would strongly depend on gap junctional connectivity across active structures. Finally, the exact nature of frequency dependence would strongly rely on the kind of active conductances expressed by the different compartments across the neuron. Given that there are no transmembrane receptor currents associated with gap junctional inputs, there is the sole reliance of extracellular potentials on the voltage-driven transmembrane currents (leak, capacitive, and active currents). Therefore, it is essential that analyses and interpretation of extracellular potentials to oscillatory inputs of different frequencies account for not just the origins of synaptic inputs, but also the membrane composition of all neurons and their dendrites in a network-specific fashion. Generalization of any kind that does not account for the specific synapses, the specific frequencies involved, or the membrane composition (including ion channels, pumps, transporters) of all cells in the network are bound to yield incorrect conclusions.

In the context of oscillations and gap-junctional coupling, electrical synapses have been shown to regulate the emergence and stability of the network interactions underlying rhythms of different frequencies, especially gamma-frequency oscillations (Draguhn et al., 1998; Hormuzdi et al., 2001; Buhl et al., 2003; LeBeau et al., 2003; Traub et al., 2003a; Konopacki et al., 2004; Bocian et al., 2009; Posluszny, 2014). Specifically, both genetic and pharmacological manipulations of gap junctions have been shown to disrupt gamma rhythms. Genetic deletion of connexin-36 impairs the gamma oscillations associated with awake, active behavioral states (Hormuzdi et al., 2001; Buhl et al., 2003). High-frequency oscillations in the hippocampus have been shown to be sensitive to pharmacological agents like carbenoxolone and octanol that are known to inhibit gap junctions. Carbenoxolone has been known to reduce the transient gamma-frequency oscillations while octanol abolishes the persistent gamma rhythm (Draguhn et al., 1998; Hormuzdi et al., 2001; Traub et al., 2003a; Posluszny, 2014). In the context of our results, where we demonstrate that the relative contributions of gap-junctional coupling to high-frequency extracellular potentials is low (Figs. 6–7), how do gap junctions contribute to enhanced extracellular gamma oscillations in these circuits?

It should be noted that in hippocampal circuits, gamma oscillations emerge predominantly due to interactions between inhibitory interneurons through GABA_A_ receptors (Whittington et al., 1995; Wang and Buzsaki, 1996; Colgin and Moser, 2010; Wang, 2010; Buzsaki and Wang, 2012; Colgin, 2016). Thus, the presence of additional gap junctional coupling between these inhibitory neurons allows for tighter synchrony between these reciprocally inhibition-coupled neurons. In other words, the presence of gap junctions increases the probability of action potential generation in other neurons that are electrically coupled to them, together increasing the population of inhibitory neurons that elicit synchronous action potentials. When these synchronous action potentials act on the adjacent cells, both excitatory and inhibitory, the transmembrane GABA_A_ receptor currents yield stronger gamma-frequency oscillations in the extracellular potentials (Draguhn et al., 1998; Hormuzdi et al., 2001; Traub et al., 2003a; Posluszny, 2014). Thus, the stronger high-frequency oscillations observed in these scenarios is owing to the enhanced synchrony that is brought about the gap-junctional coupling, which translates to stronger transmembrane inhibitory receptor currents.

These observations also strongly emphasize the utility of the computational approach we took in this study towards discerning the specific roles of gap junctions. Gap junctional coupling have strong physiological roles in terms of enhancing synchronous activity across the neurons that they couple and often express along with other receptors that connect the sets of neurons. Thus, the specific contributions of different neuronal components need to be studied with reference to how they contribute to physiological characteristics *vs.* their contributions to extracellular potentials. Thus, computational modeling offers an ideal route to understand the specific contributions of different neural-circuit components to extracellular field potentials and rhythms therein (Buzsaki et al., 2012; Einevoll et al., 2013; Einevoll et al., 2019; Sinha and Narayanan, 2022).

### Phase relationship between intracellular and extracellular potentials

The phase relationship between the intracellular voltages and extracellular potentials are dependent on several factors. First, as the receptor currents are inward and the consequent intracellular voltage response is a depolarization, that constitutes a 180° phase shift given that the receptor currents dominantly contribute to extracellular potentials associated with chemical synapses. Second, the capacitive transmembrane current for a pure oscillatory voltage response is 90° given that the capacitive current is directly related to the derivative of the voltage. The capacitive component will increase in amplitude with increasing frequencies owing to the dependence on the derivative of the voltage. Third, the presence of low-pass RC filtering implies that the voltage response is higher for low frequencies and lower at high frequencies. These differences in voltage amplitudes translate to differences in voltage-gated active channel currents, driving force differences for all transmembrane ionic currents including receptor currents, and as differences in capacitive current.

Fourth, depending on the slow and fast kinetics of the ion channels and the frequency at which the stimulation is presented, certain channel currents will be in phase with the voltage (*e.g.*, leak channels and fast outward currents), certain others will be out of phase (*e.g.*, fast calcium and sodium currents), and few others will be slow to rise and slow to fall (e.g., HCN). Thus, there will be different transmembrane currents associated with the same intracellular voltage response: some that are in phase with the voltage, some that are shifted by 90°, some at 180°, and others at arbitrary phases in between. The dominance of each of them vary with frequencies, implying that in some scenarios the capacitive current will be dominant whereas in others a specific ionic current might be dominant. Thus, the final phase of the total transmembrane current (with reference to the compartmental voltage) in each compartment would be variable depending on which component is dominant at the frequency at which the inputs arrive. Finally, as different compartments could be endowed with different densities of each ion channel, the phase of the extracellular potential recorded will be driven by the position of the recording electrodes (both extracellular and intracellular) as well. While each line current component will have a specific phase associated with the line segment being considered, the proximity of the extracellular electrode to a specific line segment as opposed to the others might enhance the contribution of that specific line segment. Similarly, as voltages within a cell also change with distance owing to RC filtering and other active filtering (Narayanan and Johnston, 2007, 2008; Vaidya and Johnston, 2013; Das et al., 2017), the position of the intracellular electrode plays a critical role in determining the phase difference between intra– and extra-cellular voltages.

With gap junctional inputs, a critical component that is absent is the 180° phase-shifted transmembrane synaptic current. The absence of this component results in strikingly different phase relationships between intra– and extra-cellular voltages in neurons receiving gap junctional inputs. Traditionally, the presence of gap junctions has been shown to enhance synchrony across neurons (Draguhn et al., 1998; Pfeuty et al., 2003). Our analyses provide evidence that the presence of gap junctional connectivity could redefine the relationship between extracellular potentials with intracellular voltages and spikes, apart from altering the polarity of extracellular signatures associated with synchronous inputs. The presence of gap junctions in oscillatory networks, such as interneuronal networks in the hippocampus that are implicated in the generation of (Posluszny, 2014) rhythms (Colgin, 2016), should be explicitly accounted for in computing and interpreting phase relationships between spikes and LFP oscillations.

### Outward HCN currents regulate extracellular potentials

An important observation from our analyses adds an intriguing facet to the multifarious physiology of HCN channels and their impact on local field potentials. There is sufficient evidence for a strong role of HCN channels in regulating local field potentials, given their subthreshold activation profile as well their slow gating kinetics (Sinha and Narayanan, 2015; Ness et al., 2016, 2018; Sinha and Narayanan, 2022). HCN channels conduct mixed cationic currents with a reversal potential of around *E_h_* = –30 mV. They are activated by hyperpolarization and their conductance value (*g*_HCN_) goes to zero at voltage values that are more depolarized than –50 mV (Robinson and Siegelbaum, 2003; Biel et al., 2009; Shah, 2014; Combe and Gasparini, 2021; Mishra and Narayanan, 2025). Thus, the current through HCN channels, *I_h_* = *g*_HCN_(*V* − *E_h_*), has always been considered to be inward (negative in sign) in nature given that *g*_HCN_ falls to zero for positive values (when *V* > *E_h_*) of the driving force (*V* − *E_h_*). In contrast to this traditional analyses associated with the polarity of HCN channels, our analyses provide two conditions under which HCN channels conduct outward currents, irrespective of whether inputs arrive through chemical synapses or gap junctions (Fig. 3; Supplementary Fig. S4).

First, the neuron should manifest a HCN conductance at resting membrane potential (RMP). Second, synaptic inputs should produce depolarizing responses that cross the reversal potential for HCN channels with fast kinetics. If these two conditions are met, then there is a duration where outward current flows through HCN channels, as a direct consequence of the slow deactivation kinetics of HCN channels. The first condition is easily met because most neurons that express HCN channels manifest a resting HCN conductance that is typically involved in regulating RMP. As RMP is typically hyperpolarized compared to the reversal potential of HCN channels, the resting current through HCN channels is inward in nature (Supplementary Fig. S4). The second requirement is achievable if neurons receive synchronous inputs that are capable of driving the membrane potentials to depolarized values beyond HCN channel reversal potential. Together, if the kinetics of the voltage response is faster than the deactivation kinetics of HCN channels, an outward current through HCN channels could manifest. This outward current could regulate intracellular neuronal computations apart from contributing to extracellular potentials as observed in our analyses. Strongly synchronous inputs are known to impinge on several neuronal subtypes expressing HCN channels (Ariav et al., 2003; Bennett and Zukin, 2004; Diba et al., 2014; Connors, 2017). Therefore, future analyses involving HCN channels should account for the possibility of outward HCN currents, both from the perspective of how they regulate cellular neurophysiology and extracellular potentials.

HCN channels play a critical role in shaping hippocampal network dynamics by modulating neuronal excitability, oscillatory behavior, and susceptibility to pathological states (Magee, 1998; Nolan et al., 2004; Kessi et al., 2022; Mishra and Narayanan, 2025). The outward-like properties of the HCN current we observed may have specific functional implications at different scales. At the cellular scale, the manifestation of outward current during action potentials or plateau potentials could contribute to afterhyperpolarization thereby regulating firing properties. In cortical and hippocampal pyramidal neurons, most single-neuron processing occurs in their elaborate dendritic branches, where there is spatiotemporal summation of different synaptic potentials, plateau potentials, backpropagating action potentials, and dendritic spikes (Johnston and Narayanan, 2008; Major et al., 2013; Stuart and Spruston, 2015). Considering the heavy expression of HCN channels in the dendrites of hippocampal and cortical pyramidal neurons (Magee, 1998; Williams and Stuart, 2000; Lorincz et al., 2002; Kole et al., 2006), the backpropagating action potentials, plateau potentials, or dendritic spikes at dendritic location could yield outward currents. These outward currents could act as a hyperpolarizing mechanism that suppresses spatiotemporal summation of the different dendritic potentials.

At the network scale, such regulation of dendritic potentials and somatic firing could contribute to overall reduction in firing rates of different neurons in the network. For instance, as inhibitory neurons typically elicit action potentials at higher frequencies, somatic outward HCN currents would occur more frequently in inhibitory neurons that express HCN channels compared to excitatory neurons. However, the heavy expression of HCN channels in the dendrites and the higher prevalence of dendritic spikes and plateau potentials in dendrites (Moore et al., 2017; Basak and Narayanan, 2018; Larkum et al., 2022) imply that the impact on outward HCN currents might be higher. Thus, the presence of outward HCN currents would regulate network balance of excitation-inhibition in an activity-dependent manner. Additionally, the outward component of the current through HCN channels could contribute to stabilization of network synchrony by promoting spike phase coherence and to modulation of spike-LFP phase relationships (Sinha and Narayanan, 2015; Ness et al., 2016; Das et al., 2017; Ness et al., 2018; Seenivasan and Narayanan, 2020; Sinha and Narayanan, 2022).

Together, the outward HCN current could play critical roles in regulating several cellular and network functions including spatiotemporal summation within single neurons, amplitude and phase of different oscillations, excitatory-inhibitory interactions, and rate and temporal coding involved in spatial navigation (O’Keefe and Recce, 1993; Nolan et al., 2004; Hussaini et al., 2011). In the context of brain rhythms, future investigations are needed to explore ripple-frequency oscillations, specifically to assess whether high-frequency network interactions are modulated by HCN outward currents. Importantly, future studies could specifically focus on delineating the prevalence and specific contributions of outward currents through HCN channels to single-neuron and network physiology.

### Limitations of analyses and future directions

Our analyses involved simulating a single morphologically and biophysically realistic neuronal model with active dendrites. We chose this approach to delineate extracellular signatures associated with the two types of synapses onto an individual neuron as well as to avoid the complexities associated with cross-interactions of signals from multiple neurons (Geisler et al., 2010; Seenivasan and Narayanan, 2020). However, LFPs are compound signals that involve several cross-interactions across different cells of various subtypes, each receiving distinct patterns of inputs through different types of synapses. Future studies should explore the implications of the stark differences between extracellular signatures associated with gap junctional *vs*. chemical synaptic inputs in networks that are endowed with different types of neurons connected through distinct synapse types.

As different neuronal subtypes are endowed with disparate sets of active dendritic components, such analyses become essential in delineating the spatio-temporal and spectral signatures of extracellular potentials associated with disparate inputs arriving at different parts of the neuron (Colgin et al., 2009; Valero et al., 2015; Fernandez-Ruiz et al., 2017; Valero and de la Prida, 2018; Navas-Olive et al., 2020; Mendoza-Halliday et al., 2024; Seenivasan et al., 2024). Such analyses become especially essential in understanding cross-*strata* or cross-laminar interactions between extracellular signals (simultaneously recorded within the same brain regions) in disparate frequency bands (Colgin et al., 2009; Valero et al., 2015; Fernandez-Ruiz et al., 2017; Valero and de la Prida, 2018; Navas-Olive et al., 2020; Mendoza-Halliday et al., 2024; Seenivasan et al., 2024). LFP analyses could be assessed in the presence of several morphologically realistic neuronal models, each receiving disparate patterns of inputs (Schomburg et al., 2012; Reimann et al., 2013; Sinha and Narayanan, 2015; Halnes et al., 2024), with details of gap junctional and chemical synaptic connectivity onto specific active dendritic structures within each subnetwork.

We demonstrated that the contribution of gap junctions to extracellular field potentials remains consistent regardless of the number of gap junctions. Specifically, we showed that the distinct positive LFP deflections persisted irrespective of their relative density on neurons (Fig. 1 with 217 gap junctions and Supplementary Fig. 1 with 99 gap junctions). Previous studies have quantitatively demonstrated that intrinsic and synaptic plasticity modulate hippocampal LFPs and phase coding (Sinha and Narayanan, 2015, 2022). Future analyses should also assess the impact of activity-dependent plasticity in ion channels (on dendrites, axonal initial segments, and other compartments), in synaptic receptors, and in gap junctions (Andersen et al., 2006; Johnston and Narayanan, 2008; Neves et al., 2008; O’Brien, 2014; Pereda, 2014; Coulon and Landisman, 2017; Magee and Grienberger, 2020; Mishra and Narayanan, 2021; Vaughn and Haas, 2022) on extracellular potentials with various kinds of gap junctional inputs and different combinations of plasticity in various structures. Interactions among different forms of plasticity and how co-dependent plasticity in different components alters extracellular field potentials could provide deeper insights about physiological changes during learning and pathological changes observed in different neurological disorders (Sinha and Narayanan, 2022).

Our analyses focused on depolarizing synaptic currents, irrespective of whether they arrived through gap junctions or chemical synapses. Further analyses could focus on hyperpolarizing synaptic currents, especially assessing scenarios where both excitation and inhibition arrive onto active structures through chemical and/or electrical synapses. Several factors, including the spatiotemporal profile of excitation and inhibition arriving onto individual neurons, the type of synapses that excitatory and inhibitory inputs recruit, balance or lack thereof between excitatory and inhibitory inputs, and electrode localization would be crucial variables in shaping extracellular potentials in these analyses. Apart from synaptic subtype and active dendritic transmembrane currents which were the focus of this study, there are several other factors, including inhomogeneities in neuropil impedance profiles and ephaptic coupling, that contribute to shaping LFPs (Buzsaki et al., 2012; Einevoll et al., 2013; Sinha and Narayanan, 2022; Halnes et al., 2024).

## METHODS

### Pyramidal neuron model

A multicompartmental, morphologically realistic, biophysical neuronal model for a CA1 pyramidal neuron was employed for all simulations. The 3D reconstruction of CA1 pyramidal neuron (*n123*) was taken from NeuroMorpho database (Ascoli et al., 2007). The models had detailed morphology with all parameters derived from electrophysiological data from pyramidal neurons of rat hippocampus. All parameters and their corresponding values for the neuronal model were derived from previously validated models (Roy and Narayanan, 2021). These CA1 models were validated against several physiological measurements along the somatodendritic axis (Roy and Narayanan, 2021). Active and passive mechanisms including individual channels, their gating kinetics, and channel distributions across the somatodendritic arbor of CA1 pyramidal neuronal models were all derived from their physiological equivalents in previous studies, as detailed below.

#### Passive mechanisms

The passive membrane properties were modeled as a combination of a capacitive current in parallel with a resistive current representing the leak channels. An axial resistivity parameter that accounted for intracellular resistance completed the passive representation of the neuronal model. Together, the parameters that defined the passive electrical properties of the neuron were: axial resistivity (*R_a_*), specific membrane resistivity (*R_m_*), and specific membrane capacitance (*C_m_*). *C_m_* and *R_a_* were set to be uniform across the neuron at 1 μF/cm^2^ and 120 τ.cm, respectively. Specific membrane resistivity (*R_m_*) varied nonuniformly in a sigmoidal manner (Golding et al., 2005; Narayanan and Johnston, 2007; Rathour and Narayanan, 2014; Basak and Narayanan, 2018) along the somatodendritic axis as a function of radial distance (*x*) of the compartments from soma:

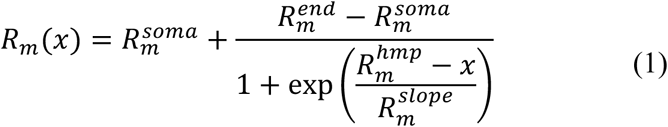

Descriptions of parameters and their default values are provided in Supplementary Table S1. The basal dendrites and axonal compartments had the same passive properties as the soma. The neuron was compartmentalized into 879 compartments using the d_λ_. rule (Carnevale and Hines, 2006), whereby each compartment was set to be smaller than 10% of λ_/00_, the space constant computed at 100 Hz. Out of these, there were 253 basal dendritic compartments.

#### Dynamics and distribution of active channels

To model the active properties of the neuron, nine different types of ion channels were incorporated into the neuron model, based on electrophysiological characterization from CA1 pyramidal neurons. Different types of ion channels were incorporated into active models: fast sodium (NaF), delayed rectifier potassium (KDR), hyperpolarization-activated cyclic nucleotide-gated non-specific cationic (HCN or *h*), *A*-type potassium (KA), *M*-type potassium (KM), *L*-type (CaL), *N*-type (CaN), *T*-type (CaT), and *R*-type (CaR) calcium channels. Current through Na, K, and HCN channels were modeled using an Ohmic formulation with the reversal potentials of Na^+^, K^+^, and HCN channels set at 55, –90, and –30 mV respectively. Current through calcium channels was modeled as per the Goldman-Hodgkin-Katz (GHK) conventions with the resting internal and external calcium concentration values set at 50 nM and 2 mM, respectively.

The conductances of NaF and KDR channels were set to be uniform across the somatodendritic arbor (Magee and Johnston, 1995; Hoffman et al., 1997). At the axonal initial segment (AIS), NaF density was tenfold higher and the conductance of KDR was double of its somatic counterpart. *N*-type calcium channels were uniformly distributed till 340 µm from soma along the apical dendrites (Magee and Johnston, 1995). *T*-type calcium channels were distributed with increasing density along the somatoapical axis as sigmoid gradient (Magee and Johnston, 1995), *M*-type potassium, and *L*-type calcium channels were distributed only at perisomatic regions till 50 µm from soma (Magee and Johnston, 1995; Hu et al., 2007). *R*-type calcium channels were distributed uniformly along somatodendritic arbor (Magee and Johnston, 1995). The gating properties and kinetics for these channels were derived from previous studies: NaF, KDR, and KA (Magee and Johnston, 1995; Hoffman et al., 1997; Migliore et al., 1999), HCN (Magee, 1998; Poolos et al., 2002), KM (Shah et al., 2008), CaT and CaN (Migliore et al., 1995), CaR and CaL (Magee and Johnston, 1995). The default conductance values associated with all ion channels are provided in Supplementary Table S1.

For KA channels, the differential kinetics and voltage-dependencies of the channel for proximal (radial distance, *x* ≤ 100 µm from the soma) and distal (*x* > 100 µm from the soma) regions were accounted for (Hoffman et al., 1997; Migliore et al., 1999). The density of KA channel was set to increase linearly (Hoffman et al., 1997) as a function of distance of the dendritic compartment from soma (Supplementary Table S1):

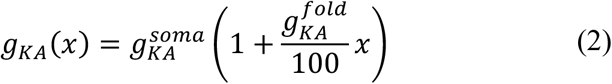

The distribution and kinetics of HCN channels were adapted from previous studies of the hippocampal recordings for pyramidal neurons (Magee, 1998; Narayanan and Johnston, 2007). The kinetics and densities of HCN channels in basal dendrites were set the same as that of soma. The gradient of maximal conductance of HCN followed a sigmoidal dependence along the radial distance, *x*, of the somatoapical arbor (Supplementary Table S1):

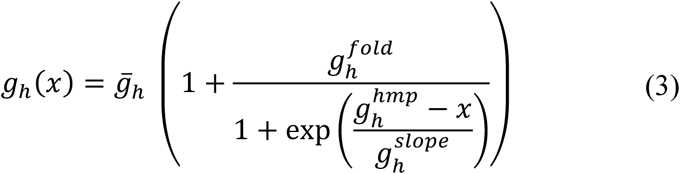

For apical dendrites, the conductance of the *T*-type calcium (CaT) channels was set to increase with radial distance as a sigmoid function (Supplementary Table S1):

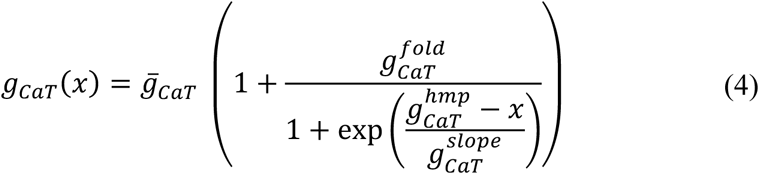

The CaT conductance for basal dendritic compartment were set to be the same as that of the soma.

These channel distributions and the associated parametric values (Supplementary Table S1) were demonstrated to satisfy 22 different somato-dendritic measurements (Roy and Narayanan, 2021). Specifically, six physiological measurements — input resistance, maximal impedance amplitude, resonance frequency, resonance strength, total inductive phase, and back-propagating action potential — were validated with respective electrophysiological ranges at three somato-dendritic locations (Soma, ∼150 µm dendrite, and ∼300 µm dendrite) each (6 × 3 = 18 measurements). In addition, action potential firing frequency at each of 100 pA, 150 pA, 200 pA, and 250 pA (4 measurements) were also matched in the model to fall within the respective ranges of corresponding electrophysiological measurements. The electrophysiological ranges of intrinsic measurements were derived from respective somato-dendritic recordings (Spruston et al., 1995; Narayanan and Johnston, 2007, 2008; Narayanan et al., 2010; Malik et al., 2016). Together, the CA1 pyramidal model neuron used in this study was tuned to match several electrophysiological characteristics and ion-channel distributions (Roy and Narayanan, 2021).

#### Models and virtual knockouts

To understand the contribution of different active dendritic transmembrane currents on extracellular potentials, we compared active (with all active conductances intact) and passive (leak channels were the only ion channels present) in the dendrites of these models. Note that in passive dendritic models, the soma continued to be active to accommodate action potential generation. For the models labeled “No sodium” and “No leak”, the sodium conductance and leak conductance were set to zero, respectively, throughout the neuron (including the soma). For the knockout model labeled “No sodium or leak”, both sodium and leak channel conductance values were set to zero across the neuron (including the soma).

### Chemical synaptic and gap junctional inputs: Characteristics and temporal dynamics

We employed excitatory chemical synapses (default number=245) which were randomly dispersed across the basal dendritic tree by picking locations from a uniform distribution of indexed locations.

The complexities of LFP modeling precludes modeling of networks of morphologically realistic models with patterns of stimulations occurring across the dendritic tree. LFP modeling studies predominantly uses post-synaptic currents to analyze the impact of different patterns of inputs arriving on to a neuron, even when chemical synaptic inputs are considered. Explicitly, individual neurons are separately simulated with different patterns of synaptic inputs, the transmembrane current at different locations recorded, and the extracellular potential is then computed using line-source approximation (Gold et al., 2006; Buzsaki et al., 2012; Schomburg et al., 2012; Reimann et al., 2013; Sinha and Narayanan, 2015; Ness et al., 2018; Sinha and Narayanan, 2022; Halnes et al., 2024). Even in scenarios where a network is analyzed, a hybrid approach involving the outputs of a point-neuron-based network being coupled to an independent morphologically realistic neuronal model is employed (Mazzoni et al., 2015; Hagen et al., 2016; Martinez-Canada et al., 2021). Given the complexities associated with the computation of electrode potentials arising as a distance-weighted summation of several transmembrane currents, these simplifications become essential.

Synaptic receptors were modeled using a two-exponential formulation (Carnevale and Hines, 2006), with rise time constant (τ_7_) and decay time constant (τ_+_) set at 5 ms and 25 ms, respectively. The reversal potential for excitatory synaptic currents was 0 mV, and the default peak conductance was set to 150 pS. The time of activation of a chemical synapse refers to the time at which the postsynaptic conductance associated with that synapse changes towards initiating the two-exponential dynamics.

Our approach models gap junctional currents in a similar way as the other model incorporate synaptic currents in LFP modeling (Gold et al., 2006; Buzsaki et al., 2012; Schomburg et al., 2012; Reimann et al., 2013; Sinha and Narayanan, 2015; Ness et al., 2018; Sinha and Narayanan, 2022; Halnes et al., 2024). As gap junctions are typically implemented as resistors from the other neuronal compartment, we accounted for gap-junctional variability in our model by randomizing the scaling-factors and the exact waveforms that arrive through individual gap junctions at specific locations from potential pre-synaptic sources. For modeling electrical synapses (Supplementary Fig. S1), membrane potential deflections from different locations were recorded from other simulated neurons (which represented the other neurons connecting through gap junctions) that received chemical synaptic inputs. The chemical synapses onto these other neurons and their dendritic distribution were modeled in similar fashion as described above. The recorded voltage waveforms from different compartments of these other neurons were differently shaped, as post-synaptic potentials or as action potentials or as dendritic spikes (Supplementary Fig. S1). These recorded voltage waveforms were normalized, converted to currents, scaled using a scaling factor, and directly injected as currents of the same sign into locations where the designated gap junctions were placed (Supplementary Fig. S1). A default number of ninety-nine randomly dispersed gap junctional inputs were placed on basal dendritic locations of the neuron under consideration. The specific waveform (from the several recorded ones) associated with each of the 99 gap junctional inputs was randomized across trials and dendritic locations. Injected current was scaled such that they generated local responses of ≤ 5 mV amplitude. Across different locations, scaling factor values ranged between 0.05–0.405 for models with active dendrites and between 0.009–0.3645 models with passive dendrites. The time of activation of a gap junction refers to the start time at which the associated current waveform was injected into the compartment where the gap junction was placed.

The precise nature of LFPs depends on spatial localization of synapses, the transmembrane currents from active structures, and on the temporal patterns of impinging activity (Buzsaki et al., 2012; Einevoll et al., 2013; Sinha and Narayanan, 2022). To assess the impact of different temporal patterns of activity on LFPs, chemical or electrical synapses were activated with three distinct temporal patterns of activity: synchronous, random, and rhythmic inputs. As the goal of this study was to assess differences in extracellular signatures associated with chemical synapses *vs*. gap junctions, the neuronal model under consideration received inputs exclusively either through chemical synapses or through gap junctions. There were no models that received both chemical synaptic as well as gap junctional inputs.

#### Synchronous synaptic inputs

Synchronous inputs were temporally synchronized excitatory inputs onto the dendrites, which are known to induce high-precision firing responses in the postsynaptic neurons (Ariav et al., 2003). All models used for the simulation with synchronous inputs were tuned to obtain only one action potential for the ease of quantitatively delineating the associated extracellular potentials. In achieving this for models with chemical synapses, the maximal sodium conductance, 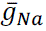 set to a default value of 30 mS/cm^2^ and maximal delayed rectifier potassium channel conductance, 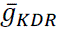 was 32 mS/cm^2^. For the models with gap junctions, the default values for 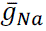 was set to 26 mS/cm^2^ and 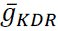 was set at 8 mS/cm^2^.

#### Random synaptic inputs

Neuronal dendrites could receive asynchronous afferents from other brain regions. We performed simulations with random asynchronous inputs that activated the neuron independently, either through chemical synapses or gap junctions. These asynchronous inputs arrived onto their targets at low– or high-frequency and were named as low-frequency random inputs (LFRI) and high-frequency random inputs (HFRI), respectively. All synapses in the group were randomly activated within a 410.5 ms window for LFRI and within a 52.45 ms window for HFRI.

#### Rhythmic synaptic inputs

Several neurons receive patterned inputs at different frequencies which yield distinct output patterns in neuronal responses (Buzsaki, 2006; Colgin, 2016). We introduced rhythmic excitatory afferent inputs at eight different frequencies: 1, 2, 4, 8, 16, 32, 64, and 128 Hz, designed to span most rhythms of the brain. Each simulation consisted of ten temporal cycles with each cycle activating 500 random locations at basal dendrites. These patterned stimuli impinged onto neurons either through chemical synapses or gap junctions. Rhythmic synaptic inputs were presented as a periodically Gaussian activation pattern with a period set at 1/*f* s and SD of 0.25/*f* s, where *f* was one of the eight frequencies above (Schomburg et al., 2012; Sinha and Narayanan, 2015).

For simulations involving these three different kinds of temporal patterns of inputs, intracellular potentials and total transmembrane currents across all compartments of the multi-compartmental neuron model were recorded and were used for further analyses.

### Positioning of extracellular electrodes and computation of extracellular potentials

We employed seven arrays of virtual electrodes, with each array made of 49 (7 × 7) electrodes, giving a total of 343 electrodes in a 7 × 7 × 7 cubic organization. These electrodes were placed over basal dendrites spanning the region from soma to the terminal end of the basal dendrites. The inter-electrode distance for each of the 3 axes was approximately 35 µm. For plotting and analysis purposes, EFP recordings from each of these electrodes were binned into 10 groups depending on their radial distance from the soma (Supplementary Table S2).

Extracellular potentials were recorded from each of these 343 electrodes as distance-scaled summation of line-source approximated currents from all line segments in the morphological realistic neuron. For each electrode, the extracellular voltage was computed as (Rall and Shepherd, 1968; Holt and Koch, 1999; Gold et al., 2006; Linden et al., 2010; Linden et al., 2011; Buzsaki et al., 2012; Schomburg et al., 2012; Einevoll et al., 2013; Linden et al., 2013; Reimann et al., 2013; Sinha and Narayanan, 2015, 2022):

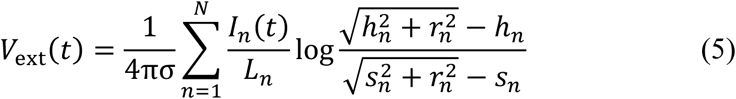

where σ defined the uniform conductivity of the extracellular medium (default value: 0.3 S/m), *I_n_* represented the line current associated with line source *n*. *L_n_* was the length of the line segment. ℎ*_n_*, *r_n_*, and *s_n_* defined geometry-related parameters that were dependent on the specific pair of electrode and line segment. *r_n_* was the perpendicular distance from the electrode to a line through the compartment, ℎ*_n_* represented the longitudinal separation along this line from the electrode to one end of the compartment, and *s_n_* was computed as the sum of ℎ*_n_* and *L*_*_. As the default integration time constant for obtaining the transmembrane currents was 25 µs, the sampling rate for *I_n_*(*t*) and *V*_ext_(*t*) was 40 kHz.

### Analysis of extracellular field potentials

Extracellular potentials recorded from all the electrodes were bandpass filtered with lower frequency cutoff of 0.5 Hz and higher frequency limit of 300 Hz to obtain local field potentials (LFPs) for that electrode. Filtering was implemented using a zero-phase, second-order, bandpass Butterworth filter on signals sampled at 40 kHz. With the synchronous stimulation paradigm, peak-to-peak field potential value were measured as the difference between peak positive and peak negative LFP deflections (within an 85-ms window after stimulation) for each electrode. The peak positive and the peak negative deflections were also computed separately for each electrode. For the random stimulation paradigm, LFPs associated with chemical *vs*. electrical stimulation were compared by computing the Fourier transform of the LFP waveform as well as the spectrogram of the LFP waveform using wavelet transform. For rhythmic synaptic inputs, each of the eight different frequency inputs (1, 2, 4, 8, 16, 32, 64, and 128 Hz) were filtered in specific bandpass frequency cut-off for higher and lower limits (Supplementary Table S3) using a zero-phase, second order (for all frequencies except for the 1 Hz, where the order was one) Butterworth filter. Peak power was obtained from the Fourier power spectrum associated with the EFP from each of the electrodes.

### Computing phase of spikes with respect to LFP oscillations associated with rhythmic synaptic inputs

For assessing spike phases associated with rhythmic synaptic inputs (at each of the 8 frequencies mentioned above), we first tuned the model using a range of synaptic weights or scaling factor for synaptic and junctional inputs, respectively, towards eliciting a single spike per oscillatory cycle. These simulations were performed over five trials, with trials differing in synaptic locations and specific inputs to these synapses. An example of parameters used in a trial of synaptic and junctional inputs are shown in Supplementary Tables S4–S5, respectively. The timing of spikes, in cycles that the neuron spiked, were recorded from soma. Extracellular potentials recorded from the electrode nearest to the soma (located 14.11 µm away from soma) was used for deriving LFPs. Extracellular potentials were filtered based on oscillatory frequency that the patterned inputs were presented through gap junctions or chemical synapses.

To assign the phase of spike in relation to LFP oscillations, the troughs from each oscillatory cycle were identified by locating the minima of field potentials for respective cycle (Seenivasan and Narayanan, 2020). The troughs were designated a phase of 0°. The phases of individual spikes were computed with reference to the troughs that preceded and followed the spike. To elaborate, let’s consider *t_spike_* as the timing of a spike elicited in response to the patterned inputs. For each spike (*t_spike_*), two troughs from the cycles were considered: one occurring just before *t_spike_*(*t_L_*), and another immediately after *t_spike_*(*t_R_*). Thus, the spike under consideration occurred between these two consecutive troughs which are separated by a phase of 360°. Therefore, phase of the spike (ϕ*_spike_*, in degrees) with reference to the oscillatory LFP was calculated as (Seenivasan and Narayanan, 2020):

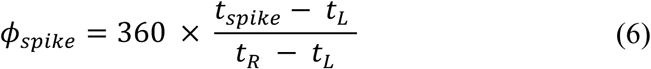

This computation was executed for all the spikes elicited by neuron in respective oscillatory cycles across all trials to assess spike phases with reference to the respective LFP oscillatory patterns.

### Computing phase of intracellular voltages with respect to LFP in models receiving rhythmic inputs

With reference to rhythmic synaptic inputs at each of the 8 frequencies mentioned above, to examine the phase of intracellular voltage traces in relation to the corresponding LFP oscillations, cross-correlation between these two waveforms was computed. To investigate the contribution of sodium conductances, we conducted cross-correlation simulations for two scenarios: one involving intracellular somatic voltages in the presence of sodium channels, filtered at respective frequencies, and another where the sodium conductance was set to zero. Parameters used for executing cross-correlation between field and somatic potentials without sodium conductance are shown in Supplementary Tables S6–S7 for synaptic and junctional inputs, respectively. Extracellular potentials were individually computed for each scenario (with sodium and no sodium) and used for cross-correlation analysis with the respective somatic voltage waveforms.

Cross-correlation was computed between the time series of soma-adjacent LFP and the associated somatic voltage. The lag (τ) corresponding to the maximum correlation was computed, and *T* (= 1⁄*f*) represented the time-period of the respective oscillation. The phase difference between the intracellular and extracellular waveforms (ϕ_DE_, in degrees) was calculated as:

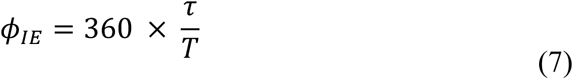

### Computational details

Morphological and biophysical properties of CA1 neuron model along with synapses, and intracellular simulations were performed using custom-written codes in the NEURON simulation environment (Carnevale and Hines, 2006). The computation of extracellular field potentials was performed using LFPy (Linden et al., 2013; Halnes et al., 2024) interfaced with NEURON (Carnevale and Hines, 2006) through Python. LFPy simulations were performed in Jupyter python terminal (https://jupyter.org) using custom-written python codes. All simulations were performed with resting membrane potential of neuron model set at –65mV and the temperature set to 34° C. The integration time step was fixed to 25 µs. Coordinates for the electrodes 3D positioning spanning dendrites were obtained using MATLAB R2015b (Mathworks). Data analyses were done using MATLAB or custom-built software written in IGOR Pro programming environment (Wavemetrics).

### Statistical analyses

All statistical analyses were performed using R statistical package (www.r-project.org). We performed Wilcoxon rank-sum test for comparing distribution of LFP data from active and passive models. In these, variability in data is shown as median with quartile. *p* values for each statistical test are provided in the figure legends associated with the specific figure panels.

## ACKNOWLEDGMENTS

The authors thank members of the cellular neurophysiology laboratory for helpful discussions and for comments on a draft of this manuscript.

## Author contributions

Conceptualization: RS, RN

Methodology: RS

Investigation: RS

Visualization: RS

Supervision: RS, RN

Writing—original draft: RS

Writing—review & editing: RS, RN

## Data and materials availability

All data are available in the main text or the supplementary materials.

## SUPPLEMENTARY INFORMATION

A supplementary file containing 12 supplementary figures and 7 supplementary tables is being uploaded.

