## Supplementary figures S1-S12; Supplementary Tables S1-S7 for "Dichotomy between extracellular signatures of active dendritic chemical synapses and gap junctions"

**This PDF file contains:**

Figures S1 to S12
Tables S1 to S7

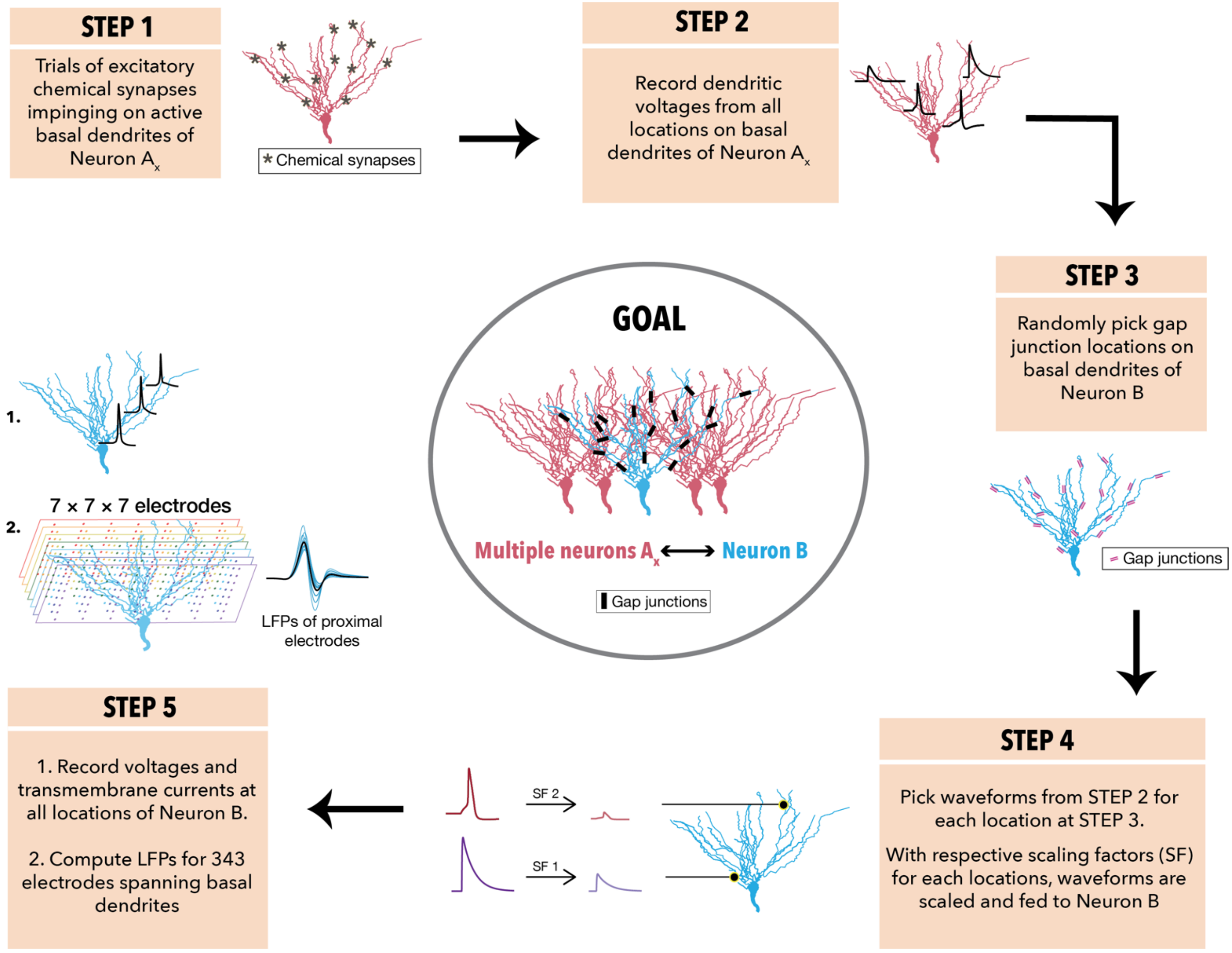


**Supplementary Figure S1. Methodology for assessing extracellular signatures of gap junctional synapses onto a neuron.** Extracellular potentials were assessed for gap junctional inputs onto the dendrites of Neuron B. To implement this, several neurons $A_x$ were considered to make gap junctional connections onto Neuron B (circle in the center named "Goal"). **Step 1**: Each neuron $A_x$ was considered to receive chemical synaptic inputs located on their active dendrites. There were no chemical synapses on Neuron B. **Step 2**: Dendritic voltages were recorded from all compartments. Depending on strength and location, these waveforms could take the form of synaptic potentials, dendritic spikes, or backpropagating action potentials. Several trials of such recordings were performed with variable localization of chemical synapses, each constituting one of the $A_x$ neuron. **Step 3**: Randomly pick gap junctional locations on Neuron B where one of the $A_x$ neurons will make contact. **Step 4**: Pick one of the waveforms from one of the trials from Step 2 and associate that waveform with one of the locations picked in Step 3. Scale the waveform (chosen from Step 2) using a location-dependent scaling factor (which represents the strength of the gap junction at the location picked in Step 3), convert the waveform into a current, and inject the current into the location chosen in Step 3. Repeat this process for all gap junctions on Neuron B. This step configures all gap junctions in Neuron B and the kind of the inputs that they receive from each of the $A_x$ neurons. **Step 5**: Record voltages and transmembrane currents from all locations of Neuron B. Use the transmembrane currents from all compartments to compute potentials at all the $7 \times 7 \times 7$ extracellular electrodes.

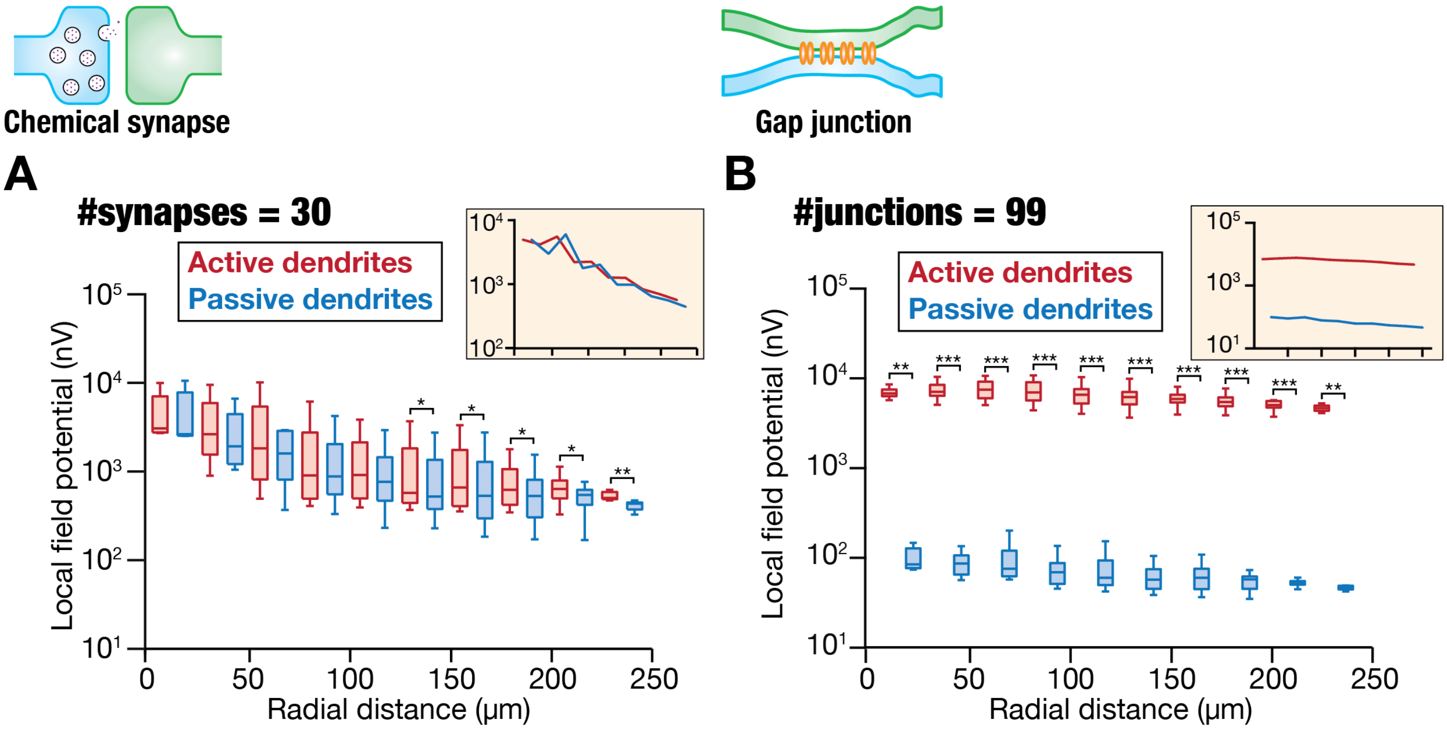


**Supplementary Figure S2. Extracellular signatures associated with active dendritic chemical synapses *vs*. gap junctions receiving synchronous inputs. *A.*** Amplitudes of negative deflection of field potentials for all 343 electrodes, plotted as functions of radial distance of the electrode from the soma, for active and passive dendritic models receiving synchronous inputs through chemical synapses ($N_{\text{syn}} = 30$). Inset shows plot of median field potential amplitude values as a function of distance for both active and passive dendritic configurations. ***B.*** Same as ***A***, but for external inputs arriving through gap junctions. The number of gap junctions $N_{\text{jun}} = 99$. Comparison of active *vs*. passive dendritic configurations in (***A–B***): *$p<0.05$, **$p<0.01$, ***$p<0.001$, Wilcoxon rank-sum test.

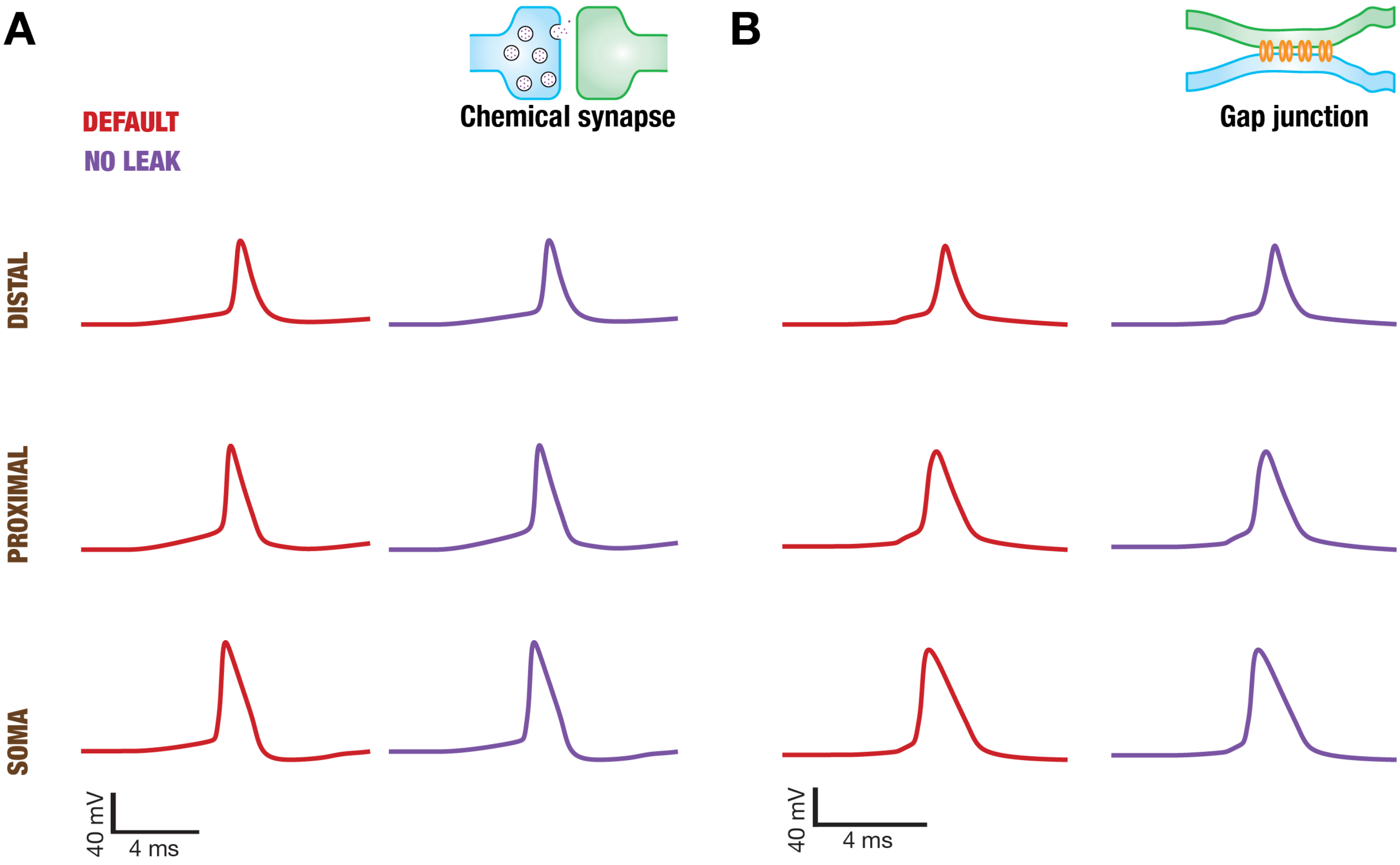


**Supplementary Figure S3. Intracellular potentials associated with active dendritic chemical synapses *vs*. gap junctions receiving synchronous inputs.** ***A***. Intracellular responses recorded at soma *(Row 1)*, proximal (100 µm; *Row 2*), and distal (200 µm; *Row 3*) dendritic locations along the somato-basal axis. Shown are the traces for default *(Column 1)* and no leak *(Column 2)* scenarios for active dendritic structures receiving synchronous inputs via chemical synapses. ***B***. same as panel *A* but with active dendrites receiving synchronous inputs through gap junctions.

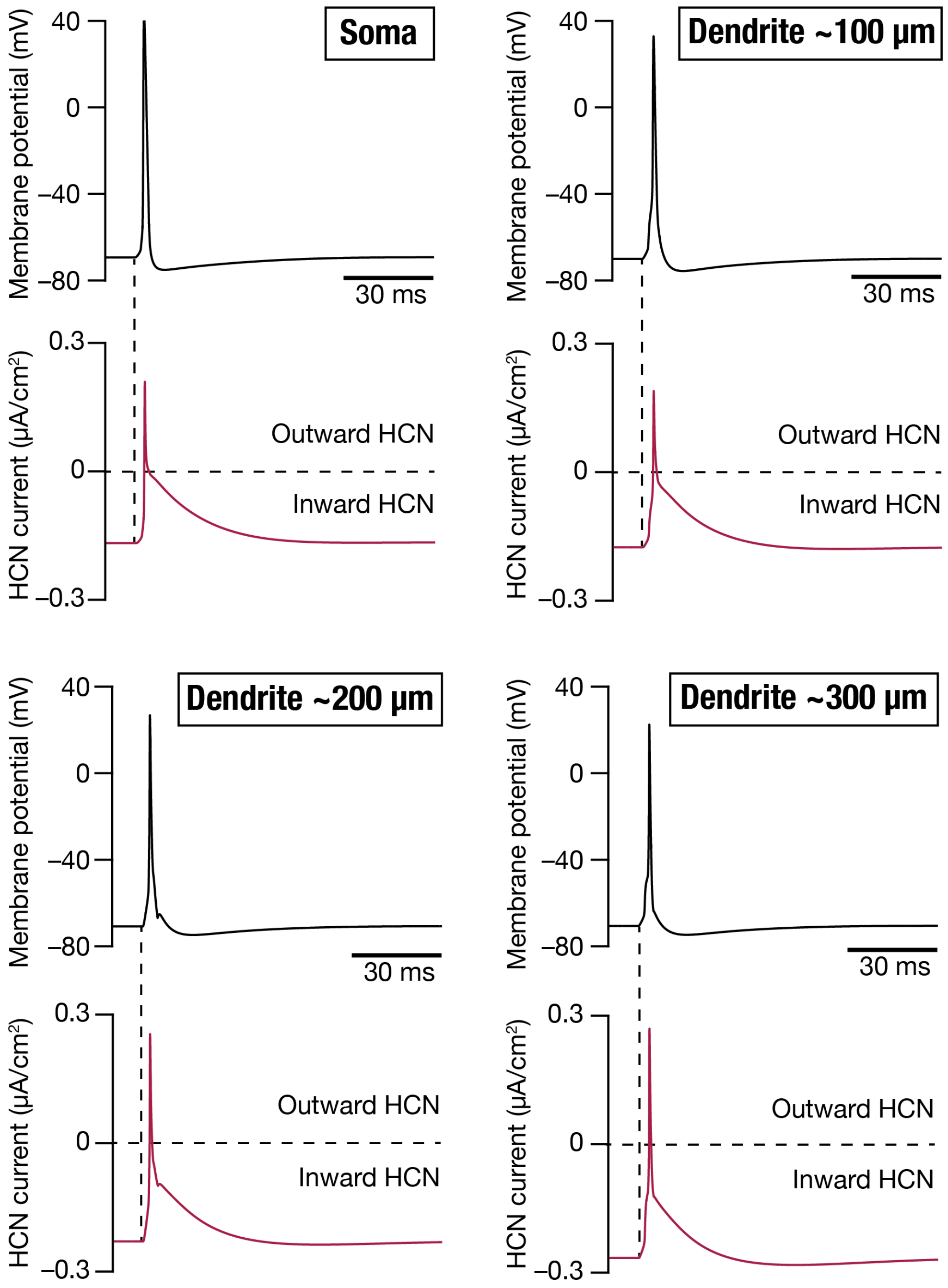


**Supplementary Figure S4. Outward transmembrane HCN currents recorded at different somatic and active dendritic compartments receiving synchronous inputs through gap junctions.** Intracellular voltages (black) and current through HCN channels (red) plotted for different somatic and dendritic (shown are distances from the soma) compartments (total duration: 110 ms). Vertical dashed lines represent the onset of synchronous inputs through gap junctions and the horizontal lines represent the transition point from inward to outward current through the HCN channels. In each case, it may be noted that under resting conditions, there is an inward (negative) resting HCN current. With the onset of synchronous stimulus, the sharp transition in voltage towards eliciting a spike alters the driving force for this resting HCN conductance. As the voltage in the compartment reaches beyond the HCN reversal potential (–30 mV), the current reverses direction to manifest an outward (positive) current through HCN channels. Thus, the resting HCN conductance combined with a sharp transition in voltage that crosses the HCN-channel reversal potential together yield an outward HCN current.

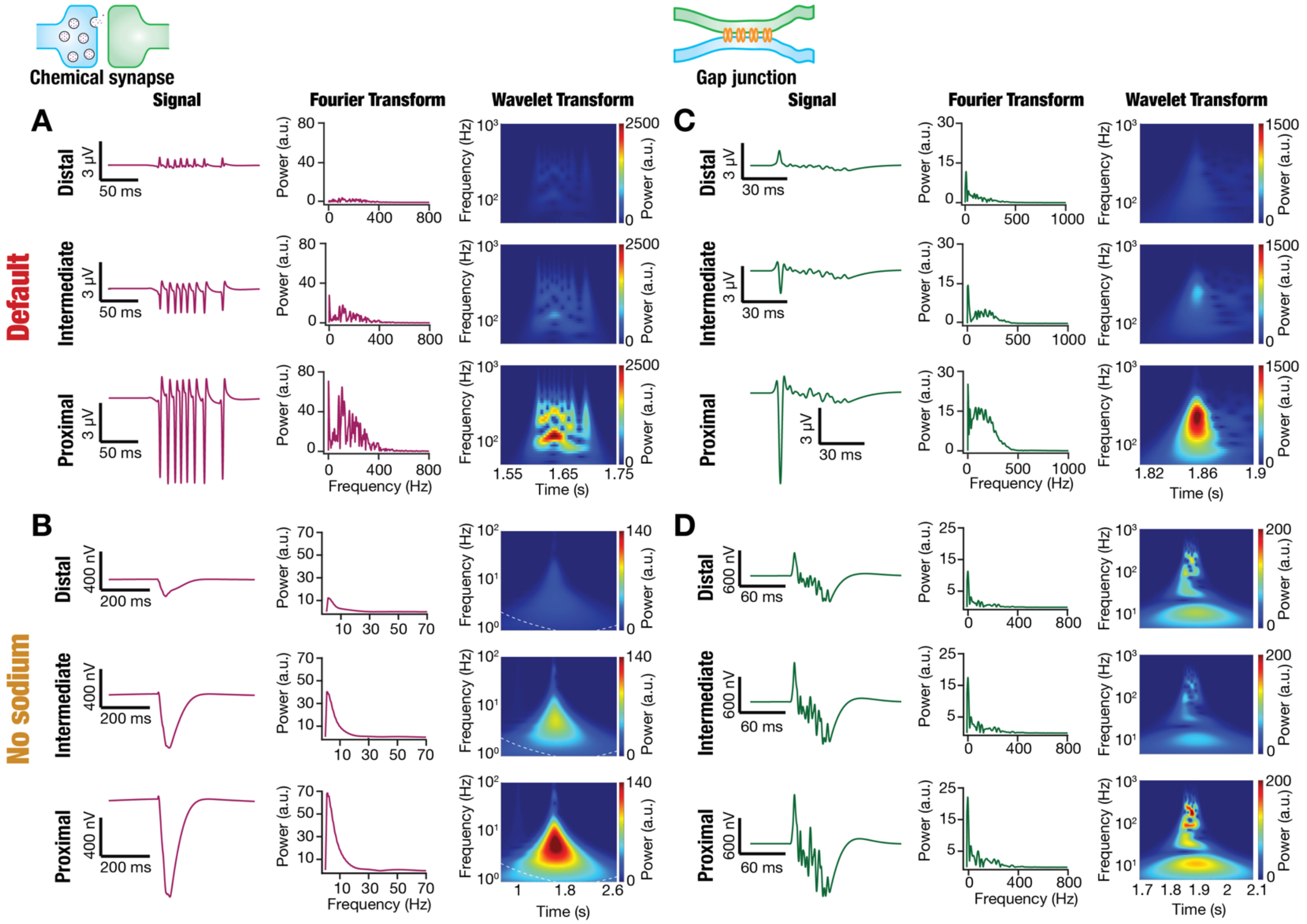


**Supplementary Figure S5**. **Differential spatiotemporal structure of field potentials associated with active dendrites receiving high-frequency random inputs through chemical synapses *vs.* gap junctions.** ***A***. Distance-wise LFP responses to high-frequency random inputs (HFRI) impinging onto active dendrites through chemical synapses. *Rows 1–3:* LFP data from electrodes located at a distal (~152 µm; *Row 1*), intermediate (~97 µm; *Row 2*), and proximal (~55 µm; *Row 3*) locations with reference to their radial distance from the soma. *Column 1:* time-domain signal. *Column 2:* Fourier transform of the signal shown in *Column 1*. *Column 3:* spectrogram of the signal shown in *Column 1* computed using wavelet transform. ***B***. Same as panel (***A***) but for active model lacking sodium conductance. ***C–D***. Same as panels (***A–B***), except high-frequency random inputs impinged onto active dendrites through gap junctions.

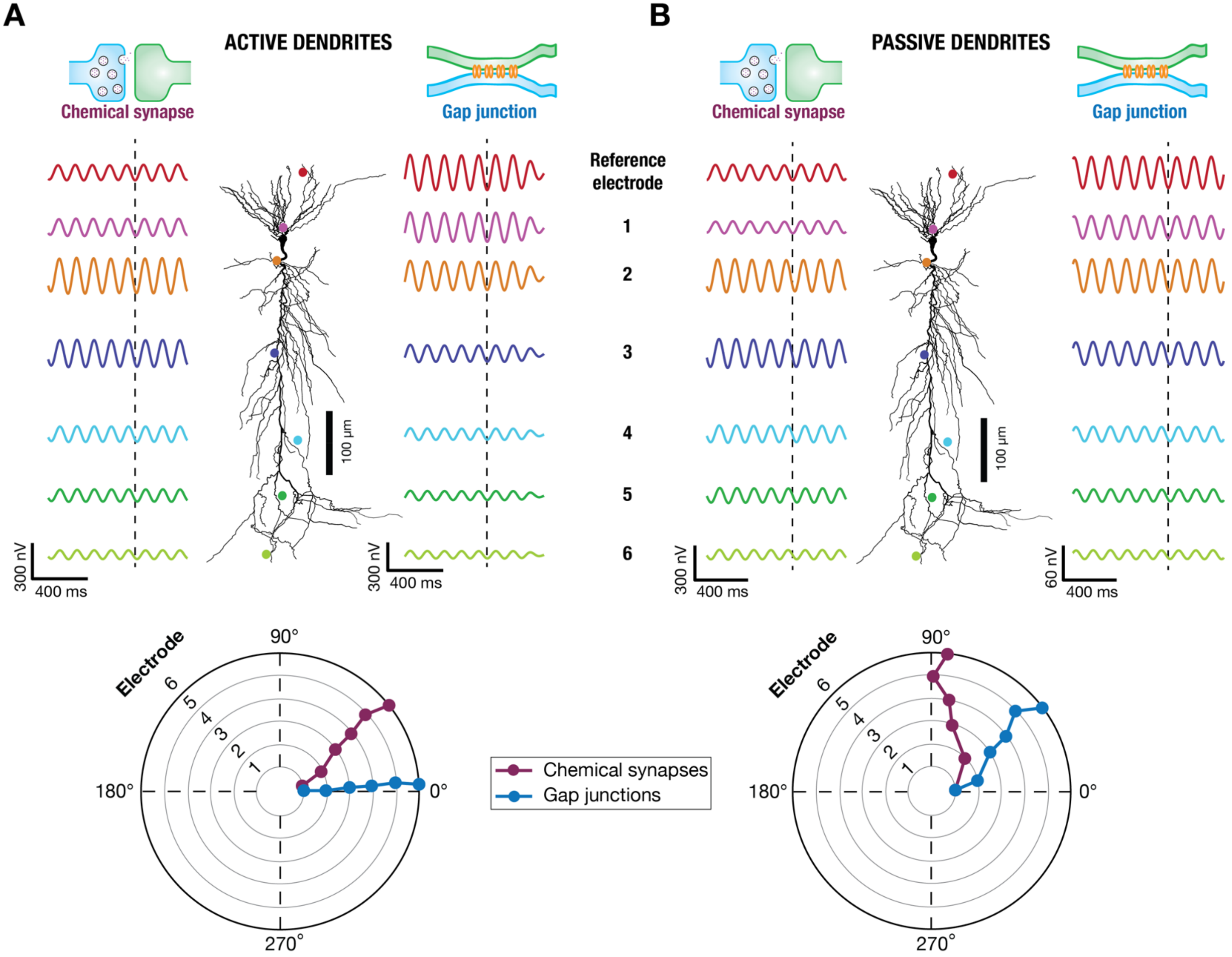


**Supplementary Figure S6**. **Somatic intracellular voltage–LFP phase relationship associated with active dendrites receiving rhythmic inputs at various frequencies through chemical synapses *vs*. gap junctions in models with no sodium channels.** ***A.*** *Top*, Extracellular potentials simultaneously recorded from electrodes that were placed across the entire span of the neuron when active dendrites received rhythmic inputs at 8 Hz through chemical synapses (*left*) or through gap junctions (*right*). 2D projections of the different electrodes and the neuronal morphology are depicted in the center. The electrode at the distal-most basal location was treated as the reference electrode to compute phase differences in the extracellular potentials observed across different electrodes. The dashed line marks a trough of the oscillatory pattern observed in the extracellular potential associated with the reference electrode. *Bottom,* Phase differences, computed through cross-correlation analysis, between the extracellular potentials recorded from reference electrode and those from electrodes 1–6, in scenarios where rhythmic inputs arrived through chemical synapses or gap junctions. ***B.*** Same as panel *A*, but for models where the dendrites were passive.

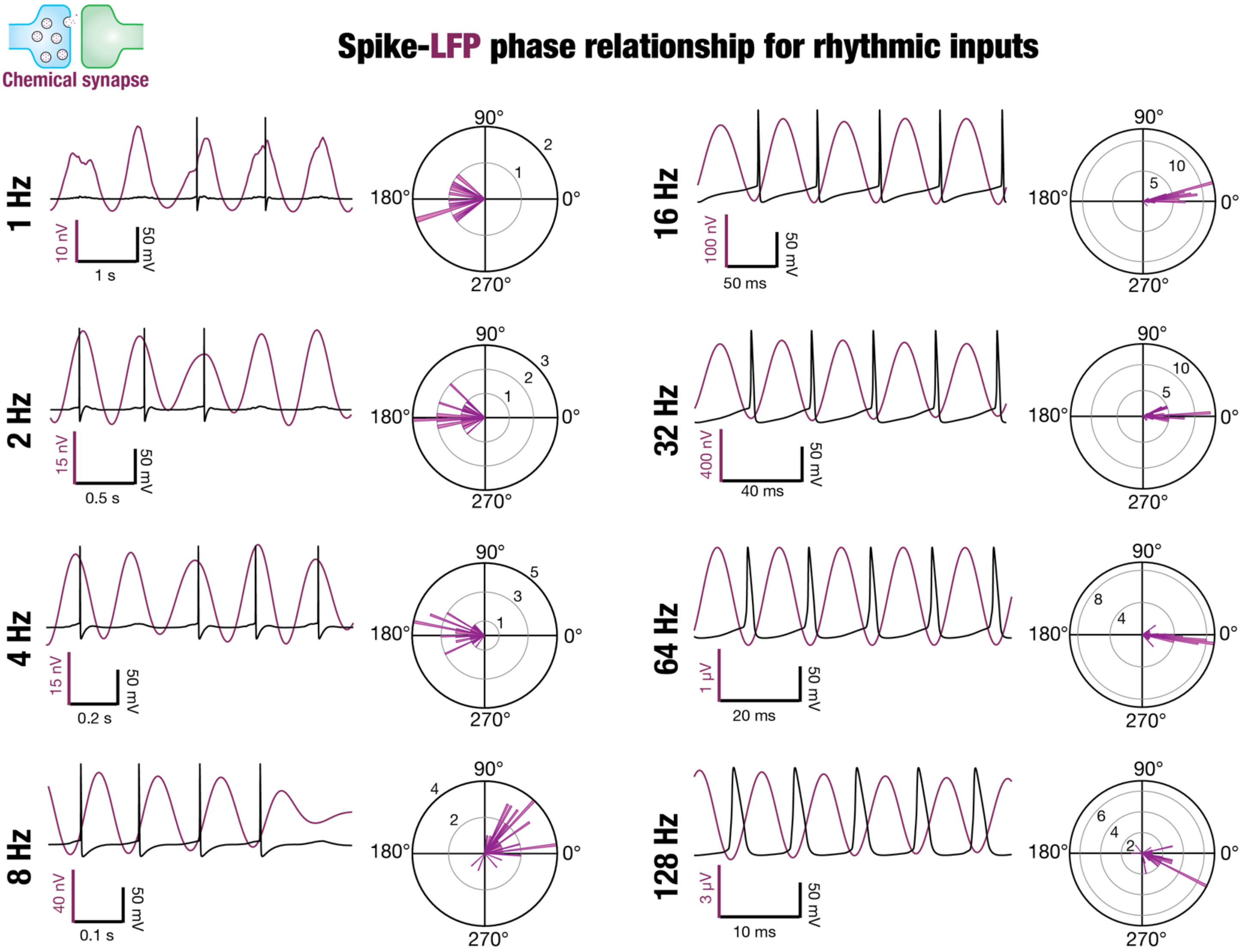


**Supplementary Figure S7**. **Somatic spike–LFP phase relationship associated with active dendrites receiving rhythmic inputs at various frequencies through chemical synapses.** Phase difference between somatic spikes and corresponding extracellular potentials at different frequencies (1–128 Hz). Note that there can be traces where spikes were not observed in all cycles. The following organization is used for each of the eight frequencies. *Column 1:* Example traces of a few cycles each of intracellular potential with spikes (black) and corresponding LFP (magenta). *Column 2:* Polar plots showing the phase difference between spikes and LFPs for different trials, with 0° indicating that the spike occurred at the trough of the LFP.

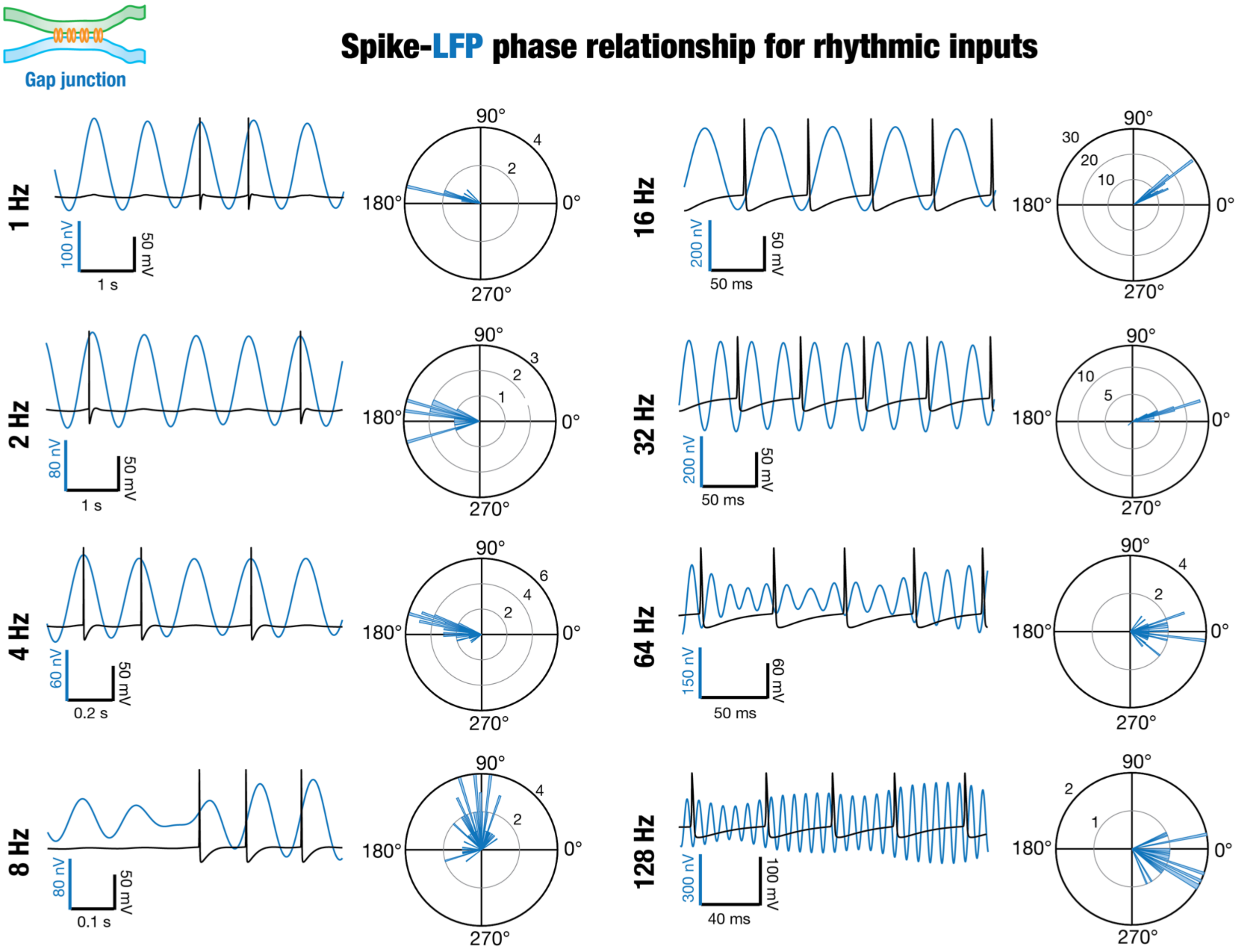


**Supplementary Figure S8**. **Somatic spike–LFP phase relationship associated with active dendrites receiving rhythmic inputs at various frequencies through gap junctions.** Phase difference between somatic spikes and corresponding extracellular potentials at different frequencies (1–128 Hz). Note that there can be traces where spikes were not observed in all cycles. The following organization is used for each of the eight frequencies. *Column 1:* Example traces of a few cycles each of intracellular potential with spikes (black) and corresponding LFP (blue). *Column 2:* Polar plots showing the phase difference between spikes and LFPs, with 0° indicating that the spike occurred at the trough of the LFP.

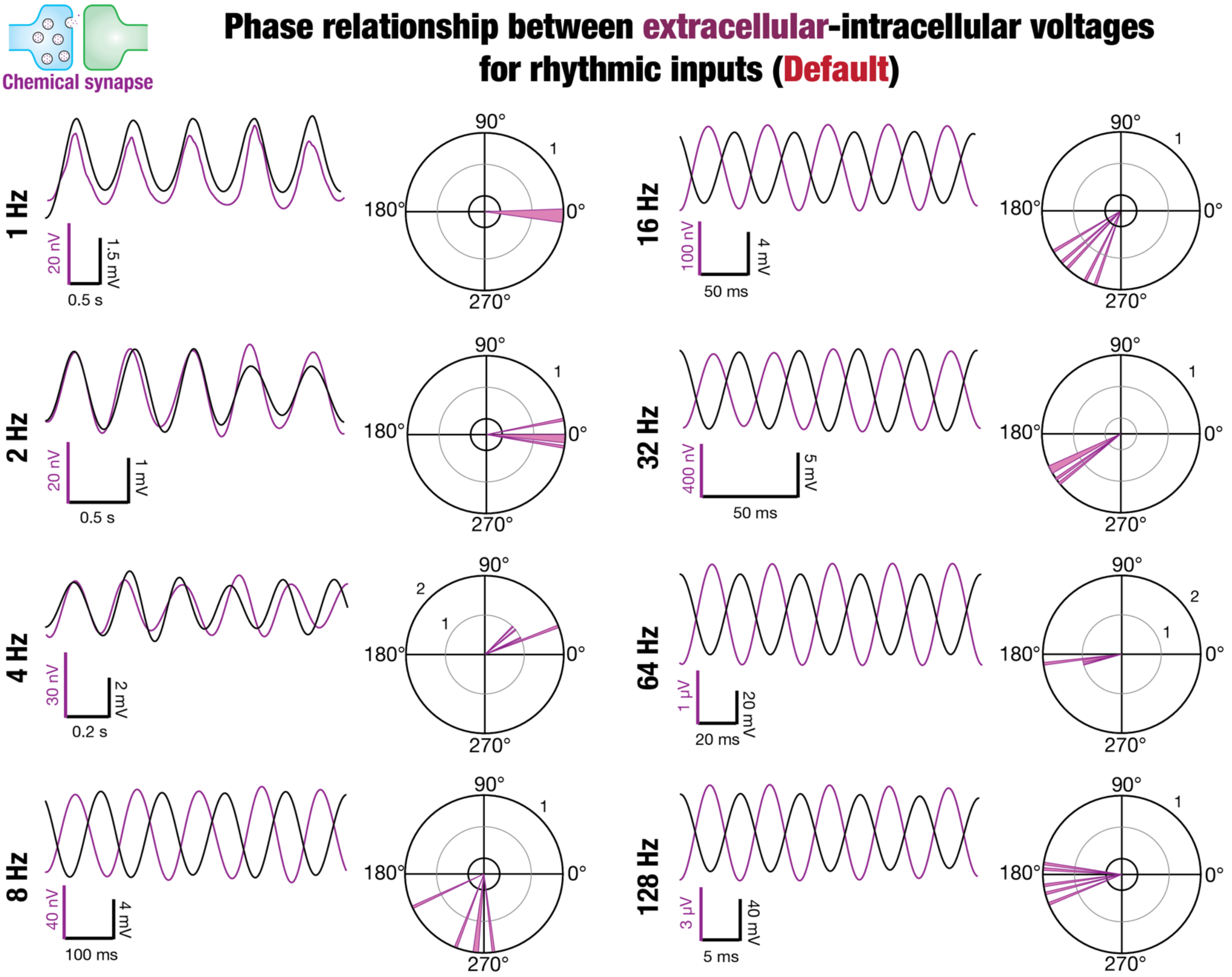


**Supplementary Figure S9**. **Somatic intracellular voltage–LFP phase relationship associated with active dendrites receiving rhythmic inputs at various frequencies through chemical synapses.** Phase difference between filtered somatic intracellular potentials (from Supplementary Fig. S6) and corresponding extracellular potentials at different frequencies (1–128 Hz). The following organization is used for each of the eight frequencies. *Column 1:* Example traces of five cycles each of intracellular potential (black) and corresponding LFP (magenta). *Column 2:* Polar plots showing the phase difference between filtered intracellular recordings and LFPs, with 0° indicating in-phase relationship between intracellular voltage and LFP.

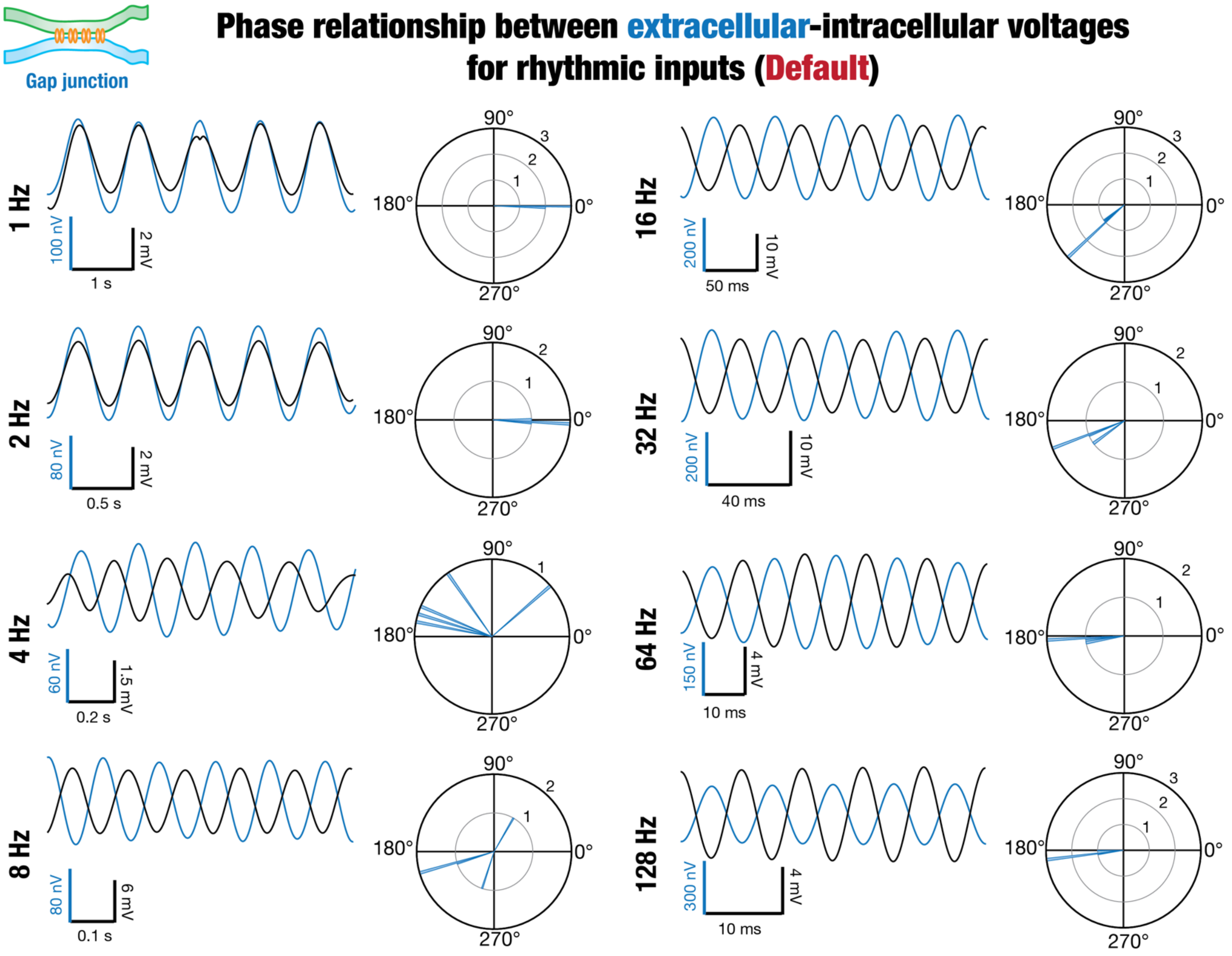


**Supplementary Figure S10**. **Somatic intracellular voltage–LFP phase relationship associated with active dendrites receiving rhythmic inputs at various frequencies through gap junctions.** Phase difference between filtered somatic intracellular potentials (from Supplementary Fig. S7) and corresponding extracellular potentials at different frequencies (1–128 Hz). The following organization is used for each of the eight frequencies. *Column 1:* Example traces of five cycles each of intracellular potential (black) and corresponding LFP (blue). *Column 2:* Polar plots showing the phase difference between filtered intracellular recordings and LFPs, with 0° indicating in-phase relationship between intracellular voltage and LFP.

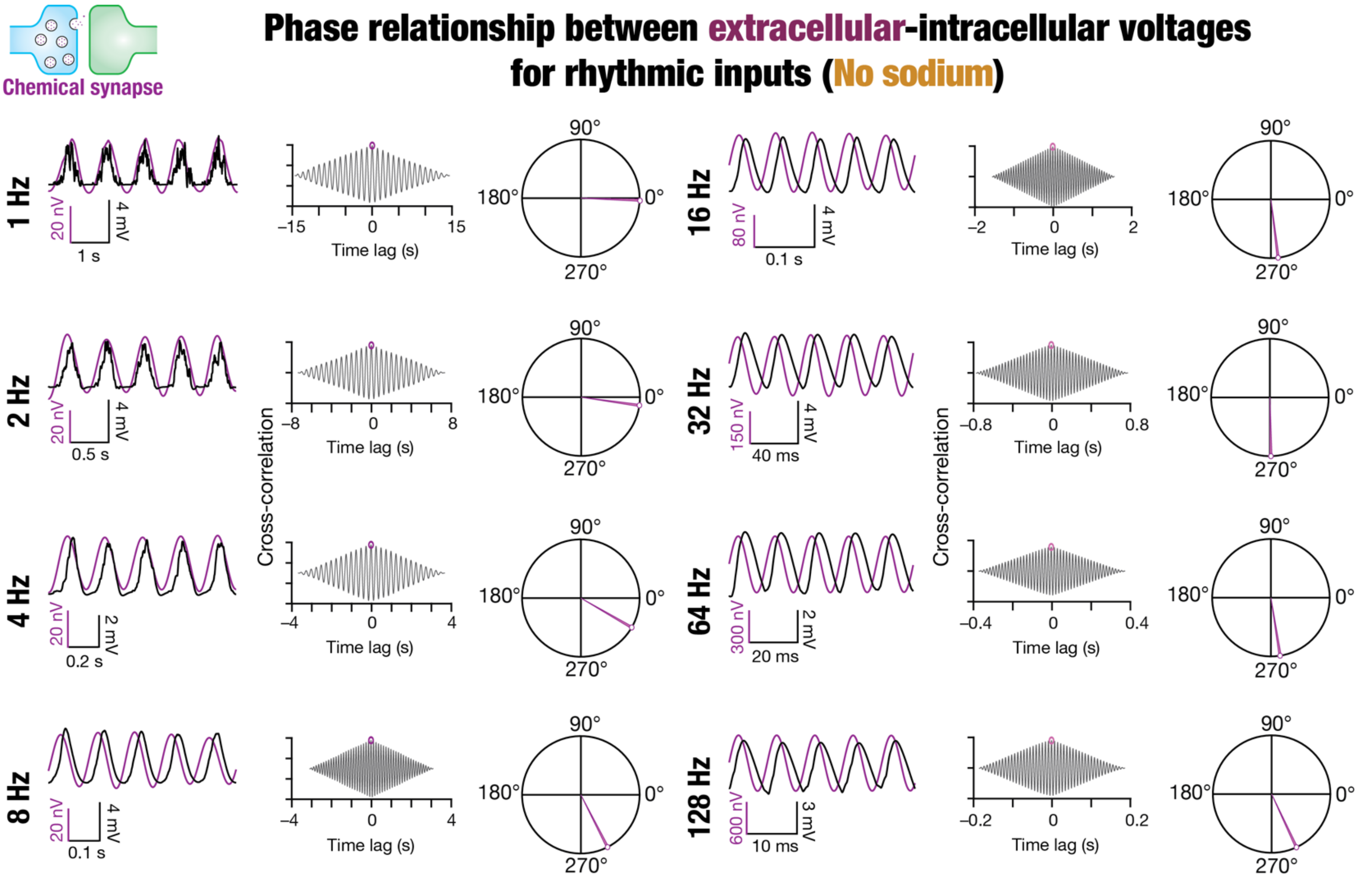


**Supplementary Figure S11**. **Somatic intracellular voltage–LFP phase relationship associated with active dendrites receiving rhythmic inputs at various frequencies through chemical synapses in models with no sodium channels.** Traces and plots showing phase difference between somatic intracellular potentials (in the absence of sodium conductance) and corresponding extracellular potentials at different frequencies (1–128 Hz). The following organization is used for each of the eight frequencies. *Column 1:* Example traces of 5 cycles each of intracellular potential (black) and corresponding LFP (magenta). *Column 2:* Cross-correlogram between these waveforms, highlighting the peak value with a magenta circle. *Column 3:* Polar plots showing the phase difference between intracellular recordings and LFPs, with 0° indicating in-phase relationship between intracellular voltage and LFP.

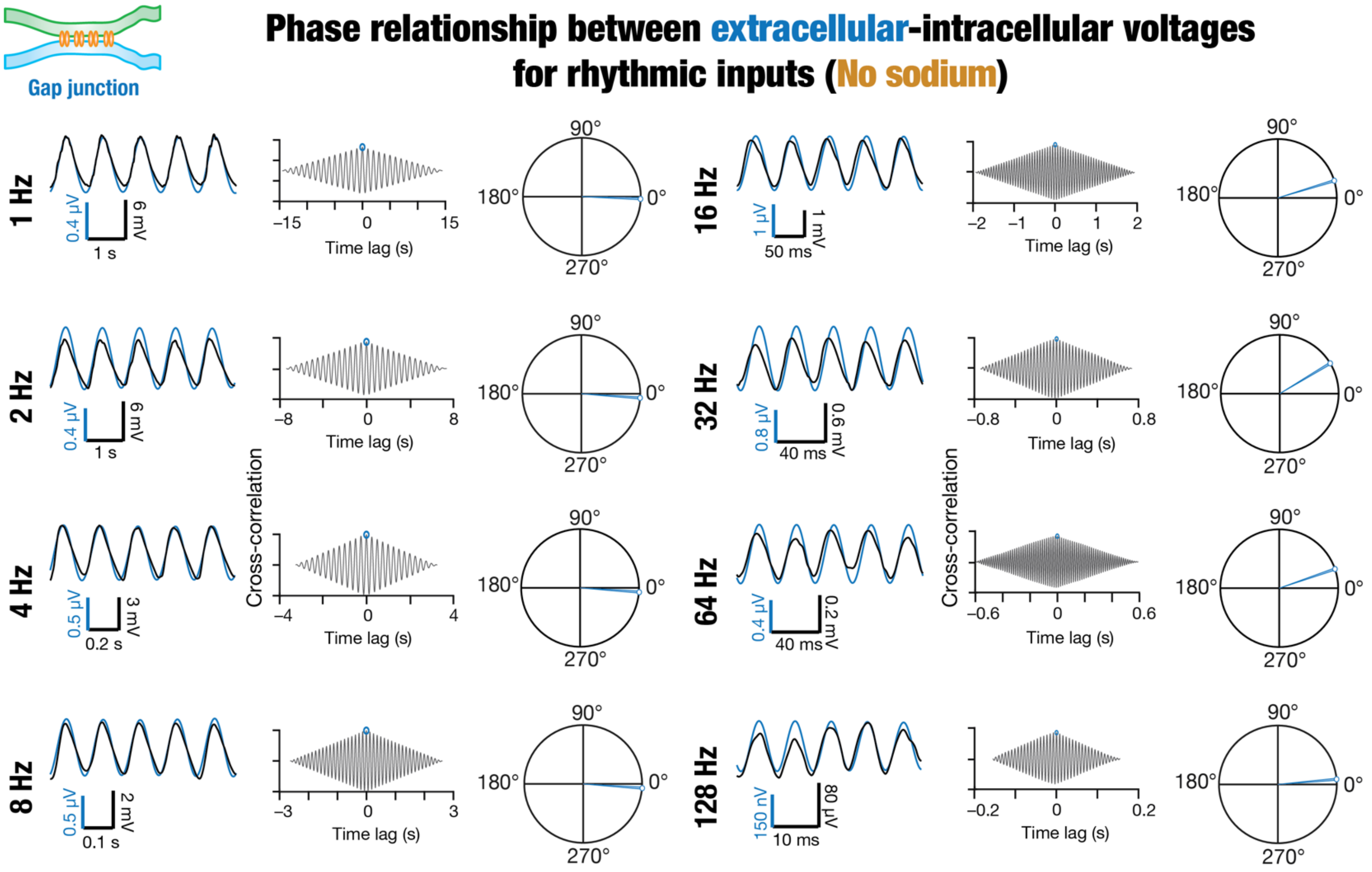


**Supplementary Figure S12**. **Somatic intracellular voltage–LFP phase relationship associated with active dendrites receiving rhythmic inputs at various frequencies through gap junctions in models with no sodium channels.** Traces and plots showing phase difference between somatic intracellular potentials (in the absence of sodium conductance) and corresponding extracellular potentials at different frequencies (1–128 Hz). The following organization is used for each of the eight frequencies. *Column 1:* Example traces of 5 cycles each of intracellular potential (black) and corresponding LFP (blue). *Column 2:* Cross-correlogram between these waveforms, highlighting the peak value with a blue circle. *Column 3:* Polar plots showing the phase difference between intracellular recordings and LFPs, with 0° indicating in-phase relationship between intracellular voltage and LFP.

**Supplementary Table S1**: Biophysical simulation parameters. These parameters were derived from (Roy and Narayanan, 2021).

| Parameter (gradient) | Unit | Symbol | Value |
|---|---|---|---|
| ***Passive properties*** | | | |
| 1 Axial resistivity<br>(Uniform across the neuron) | Ω.cm | $R_a$ | 120 |
| 2 Specific membrane resistivity<br>(Sigmoid reduction along radial distance)<br>Somatic value<br>Smallest value<br>Slope of sigmoid<br>Half-maximal point of sigmoid | <br><br>kΩ.cm$^2$<br>kΩ.cm$^2$<br>µm<br>µm | $R_m$<br><br>$R_m^{soma}$<br>$R_m^{end}$<br>$R_m^{slope}$<br>$R_m^{hmp}$ | <br><br>120<br>85<br>50<br>300 |
| 3 Specific membrane capacitance | µF/cm$^2$ | $C_m$ | 1 |
| ***Active properties*** | | | |
| 4 Spike generating channels<br>(Uniform across somatodendritic axis)<br>Default maximal conductance of NaF<br>Default maximal conductance of KDR | <br><br>mS/cm$^2$<br>mS/cm$^2$ | <br><br>$\bar{g}_{Na}$<br>$\bar{g}_{KDR}$ | <br><br>26<br>8 |
| 5 HCN<br>(Sigmoidal increase along radial distance)<br>Maximal somatic conductance of HCN<br>Fold increase<br>Slope of sigmoid<br>Half-maximal point of sigmoid | <br><br>µS/cm$^2$<br>—<br>µm<br>µm | <br><br>$\bar{g}_h$<br>$g_h^{fold}$<br>$g_h^{slope}$<br>$g_h^{hmp}$ | <br><br>25<br>12<br>50<br>320 |
| 6 *A*-type potassium, KA<br>(Linear increase along radial distance)<br>Maximal conductance of KA<br>Fold increase per 100 µm | <br><br>mS/cm$^2$<br>— | <br><br>$\bar{g}_{KA}$<br>$g_{KA}^{fold}$ | <br><br>3.1<br>8 |
| 7 *M*-type potassium, KM<br>(Perisomatic; till 50 µm from soma)<br>Maximal conductance of KM | <br><br>µS/cm$^2$ | <br><br>$\bar{g}_{KM}$ | <br><br>1 |
| 8 *T*-type calcium, CaT<br>(Sigmoidal increase along radial distance)<br>Maximal conductance of CaT<br>Fold increase<br>Slope of sigmoid<br>Half-maximal point of sigmoid | <br><br>µS/cm$^2$<br><br>µm<br>µm | <br><br>$\bar{g}_{CaT}$<br>$g_{CaT}^{fold}$<br>$g_{CaT}^{slope}$<br>$g_{CaT}^{hmp}$ | <br><br>80<br>30<br>50<br>350 |
| 9 *N*-type calcium, CaN<br>Maximal conductance of CaN | <br>µS/cm$^2$ | <br>$\bar{g}_{CaN}$ | <br>15 |
| 10 *R*-type calcium, CaR<br>Maximal conductance of CaR | <br>µS/cm$^2$ | <br>$\bar{g}_{CaR}$ | <br>15 |
| 11 *L*-type calcium, CaL<br>Maximal conductance of CaL | <br>mS/cm$^2$ | <br>$\bar{g}_{CaL}$ | <br>1.2 |

**Supplementary Table S2:** Distance ranges associated with bins for extracellular and intracellular plots.

| Bins | Range of radial distances (μm) |
|---|---|
| | ***Extracellular*** |
| **1** | 14.12 – 35.85 |
| **2** | 35.85 – 57.58 |
| **3** | 57.58 – 79.31 |
| **4** | 79.31 – 101.05 |
| **5** | 101.05 – 122.78 |
| **6** | 122.78 – 144.51 |
| **7** | 144.51 – 166.25 |
| **8** | 166.25 – 187.98 |
| **9** | 187.98 – 209.71 |
| **10** | 209.71 – 231.44 |
| | ***Intracellular*** |
| **1** | 0 – 50 |
| **2** | 50 – 100 |
| **3** | 100 – 150 |
| **4** | 150 – 200 |
| **5** | 200 – 300 |

**Supplementary Table S3:** Bandpass filtering frequency ranges for analyzing extracellular potentials associated with rhythmic inputs.

| Frequency of rhythmic inputs (Hz) | Lower limit cut-off (Hz) | Upper limit cut-off (Hz) |
|---|---|---|
| **1** | 0.5 | 1.5 |
| **2** | 1.5 | 3 |
| **4** | 3 | 5 |
| **8** | 6 | 10 |
| **16** | 10 | 20 |
| **32** | 30 | 40 |
| **64** | 60 | 70 |
| **128** | 120 | 132 |

**Supplementary Table S4:** Parameters, associated with one of the trials, used in spike-LFP phase for synaptic inputs. Frequencies in the range of 1–8 Hz were classified as low frequencies, whereas 16–128 Hz were classified as high frequencies.

| Frequency (Hz) | Duration (s) | # cycles | Weight (pS) | # spikes |
|---|---|---|---|---|
| **1** | 15 | 15 | 200 | 4 |
| **2** | 8 | 15 | 143.3 | 7 |
| **4** | 4.5 | 15 | 91 | 13 |
| **8** | 4 | 25 | 53.1 | 22 |
| **16** | 2.5 | 25 | 42 | 25 |
| **32** | 2.5 | 25 | 42 | 25 |
| **64** | 1.5 | 25 | 58.5 | 25 |
| **128** | 1.5 | 25 | 75.82 | 25 |

**Supplementary Table S5:** Parameters used in spike-LFP phase for junctional inputs in an example trial. Frequencies in the range of 1–8 Hz were classified as low frequencies, whereas 16–128 Hz were classified as high frequencies.

| Frequency (Hz) | Duration (s) | # cycles | Scaling factor | # spikes |
|---|---|---|---|---|
| **1** | 15 | 15 | 0.403 | 8 |
| **2** | 8 | 15 | 0.27173 | 2 |
| **4** | 4.5 | 15 | 0.174 | 4 |
| **8** | 4 | 25 | 0.108 | 21 |
| **16** | 3.5 | 40 | 0.060 | 35 |
| **32** | 2.2 | 40 | 0.029 | 9 |
| **64** | 2 | 45 | 0.0168 | 7 |
| **128** | 1.6 | 45 | 0.012 | 4 |

**Supplementary Table S6:** Parameters used in cross-correlation for synaptic inputs. Frequencies in the range of 1–8 Hz were classified as low frequencies, whereas 16–128 Hz were classified as high frequencies.

| Frequency (Hz) | Duration (s) | # cycles | Weight (pS) |
|---|---|---|---|
| **1** | 15 | 15 | 250 |
| **2** | 8 | 15 | 150 |
| **4** | 4.5 | 15 | 58 |
| **8** | 4 | 25 | 40 |
| **16** | 2.5 | 25 | 42 |
| **32** | 2.5 | 25 | 55 |
| **64** | 1.5 | 25 | 92 |
| **128** | 1.5 | 25 | 460 |

**Supplementary Table S7:** Parameters used in cross-correlation for gap junctional inputs. Frequencies in the range of 1–8 Hz were classified as low frequencies, whereas 16–128 Hz were classified as high frequencies.

| Frequency (Hz) | Duration (s) | # cycles | Scaling factor |
|---|---|---|---|
| **1** | 15 | 15 | 1.25 |
| **2** | 8 | 15 | 1.25 |
| **4** | 4.5 | 15 | 1.25 |
| **8** | 4 | 25 | 1.25 |
| **16** | 3.5 | 40 | 3 |
| **32** | 2.2 | 40 | 5.2 |
| **64** | 2.5 | 110 | 5.2 |
| **128** | 1.6 | 110 | 5 |
